# A post-transcriptional regulatory code for mRNA stability during the zebrafish maternal-to-zygotic transition

**DOI:** 10.1101/292441

**Authors:** Charles E. Vejnar, Mario Abdel Messih, Carter M. Takacs, Valeria Yartseva, Panos Oikonomou, Romain Christiano, Marlon Stoeckius, Stephanie Lau, Miler T. Lee, Jean-Denis Beaudoin, Hiba Darwich-Codore, Tobias C. Walther, Saeed Tavazoie, Daniel Ci- fuentes, Antonio J. Giraldez

**Affiliations:** Department of Genetics, Yale University School of Medicine, New Haven, CT 06510 USA; Yale Stem Cell Center, Yale University School of Medicine, New Haven, CT 06510 USA; Yale Cancer Center, Yale University School of Medicine, New Haven, CT 06510 USA; University of New Haven, West Haven, CT 06516 USA; Department of Molecular Biology, Genentech, Inc. South San Francisco, CA 94080 USA; Department of Systems Biology, Columbia University, New York, NY 10032 USA; Department of Genetics and Complex Diseases, Harvard T.H. Chan School of Public Health, Boston, MA 02115 USA; New York Genome Center, New York, NY 10013 USA; Department of Biological Sciences, University of Pittsburgh, Pittsburgh, PA 15260 USA; Department of Cell Biology, Harvard Medical School, Boston, MA 02115 USA; Broad Institute of Harvard and MIT, Cambridge, MA 02124 USA; Howard Hughes Medical Institute, Boston, MA 02115 USA; Department of Biochemistry and Molecular Biophysics, and Department of Systems Biology, Columbia University, New York, NY 10032 USA; Department of Biochemistry, Boston University School of Medicine, Boston, MA 02118

## Abstract

Post-transcriptional regulation is crucial to shape gene expression. During the Maternal-to-Zygotic Transition (MZT), thousands of maternal transcripts are regulated upon fertilization and genome activation. Transcript stability can be influenced by *cis-*elements and *trans-*factors, but how these inputs are integrated to determine the overall mRNA stability is unclear. Here, we show that most transcripts are under combinatorial regulation by multiple decay pathways during zebrafish MZT. To identify *cis*-regulatory elements, we performed a massively parallel reporter assay for stability-influencing sequences, which revealed that 3’-UTR poly-U motifs are associated with mRNA stability. In contrast, miR-430 target sequences, UAUUUAUU AU-rich elements (ARE), CCUC and CUGC elements emerged as the main destabilizing motifs in the embryo, with miR-430 and AREs causing mRNA deadenylation in a genome activation-dependent manner. To identify the *trans-*factors interacting with these *cis-*elements, we comprehensively profiled RNA-protein interactions and their associated regulatory activities across the transcriptome during the MZT. We find that poly-U binding proteins are preferentially associated with 3’-UTR sequences and stabilizing motifs, and that antagonistic sequence contexts for poly-C and poly-U binding proteins shape the binding landscape and magnitude of regulation across the transcriptome. Finally, we integrate these regulatory motifs into a machine learning model that accurately predicts the stability of mRNA reporters *in vivo*. Our findings reveal how mechanisms of post-transcriptional regulation are coordinated to direct changes in mRNA stability within the early zebrafish embryo.

## Introduction

Post-transcriptional regulation plays an essential role in shaping gene expression. 3’-UTRs represent a central regulatory hub that integrate multiple inputs to the mRNA, regulating its translation, localization, stability and polyadenylation status (Mayr, 2017). In the cell, this regulatory input through the 3’-UTR integrates signals from RNA binding proteins and non-coding sequences such as microRNA binding sites and AU-rich elements. Together with codon optimality and RNA modifications, they regulate mRNA stability in the cell (Schoenberg et al, 2012; Gilbert et al, 2016; Hanson et al, 2017; Mayr, 2017). The search for non-coding regulatory elements has focused on the sequence of individual 3’-UTRs (Voeltz et al, 1998; Wirsing et al, 2011; Kristjánsdóttir et al, 2015). Transcriptome-wide analysis of mRNA stability, using pulse labeling (Miller et al, 2011) or blocking transcription (Geisberg et al, 2014), have led to the identification of potential regulatory sequences by searching for common motifs within mRNA. Parallel reporter libraries (Zhao et al, 2014; Oikonomou et al, 2014; Wissink et al, 2016; Rabani et al, 2017) have been used to identify the regulatory sequences within 3’-UTRs. However, these approaches have been limited in their sequence diversity and length. For example, short sequences can be limiting when defining how the sequence context influences regulation by individual motifs, while longer sequences lack resolution. To address these issues, we recently developed a high-throughput RNA-element selection assay (RESA) to measure *in vivo* the regulatory activities of mRNA sequences (Yartseva et al, 2017). RESA uses endogenous RNA fragments in a parallel reporter assay that allows high sequence complexity and high density coverage of the transcriptome, or targeted regions of interest, providing near nucleotide resolution of the regulatory activity of RNA sequences.

RNA regulatory elements are recognized by *trans*-factors, including miRNAs and RNA-binding proteins (RBPs) (Glisovic et al, 2008). RBPs can regulate the processing, stability and translation of their target mRNAs (Gerstberger et al, 2014). Recent approaches have revealed a set of proteins in intimate contact with mRNAs across different eukaryotic systems, through interactome capture (Castello et al, 2012; Kwon et al, 2013; Wessels et al, 2016; Sysoev et al, 2016; Despic et al, 2017). To define targeting elements within RNA, *in vitro* affinity selection methods, such as SELEX (Tuerk et al, 1990; Blackwell et al, 1990; Ellington et al, 1990), RNA affinity profiling (Tome et al, 2014) and RNAcompete (Ray et al, 2013), have been complemented with UV cross linking and immunoprecipitation to provide the set of targets and the binding motifs *in vivo* for a number of RBPs (Ule et al, 2003; van der Brug et al, 2008; Chi et al, 2009; Hafner et al, 2010; König et al, 2010; Chan et al, 2014; Hansen et al, 2015; Murn et al, 2015; Sugimoto et al, 2015; Galloway et al, 2016; Scheckel et al, 2016; Rot et al, 2017). However, the presence of a specific sequence motif is not always indicative of regulation *in vivo*, suggesting that additional sequences, or combinatorial interactions between RBPs influence the regulatory output on an mRNA. Current efforts have not yet linked the RNA regulatory maps with the RBP binding profile to define the post-transcriptional regulatory network *in vivo*.

Identifying functional regulatory sequences together with RBP *trans*-factors is an essential step toward understanding post-transcriptional regulatory code of the cell. Post-transcriptional regulation is of particular importance during the early stages of animal development, which are driven by maternally provided mRNAs (Walser et al, 2011). During the Maternal-to-Zygotic Transition (MZT), mRNAs deposited in the oocyte undergo coordinated remodeling. Individual pathways have been implicated in the regulation of maternal mRNAs (reviewed in Lee et al, 2014; Yartseva et al, 2015). For example, in *Drosophila*, the RBP SMAUG destabilizes maternal mRNAs (Dahanukar et al, 1999; Tadros et al, 2007). In *Xenopus*, AU-rich elements (ARE) within the 3’-UTRs of maternal mRNAs trigger their deadenylation after egg activation, and decay after the mid-blastula transition (MBT) (Audic et al, 1997; Voeltz et al, 1998). In zebrafish, zygotic transcription of microRNA miR-430 regulates ~20% of destabilized maternal transcripts (Giraldez et al, 2006). Codon usage in the coding sequence of mRNAs predicts differential mRNA stability during the MZT across several vertebrates (Bazzini et al, 2016; Mishima et al, 2016), and it influences mRNA half-life in yeast (Presnyak et al, 2015). mRNA methylation has been implicated in shaping mRNA stability during ES cell differentiation (Batista et al, 2014) and the MZT (Zhao et al, 2017). Despite these singular discoveries, there is a lack of quantitative models that integrate the various elements to predict mRNA deadenylation and degradation. Thus, it is still poorly understood how overall sequence composition influences mRNA stability, how the sequence environment affects the regulatory potential of each motif, what relative activities different elements contribute within the same RNA and which RBPs mediate this regulation.

To decipher the post-transcriptional regulatory code shaping mRNA stability during embryogenesis, we set out to identify the *cis*-regulatory sequences and the associated *trans*-factor RBPs, with the goal to develop a quantitative model that explain the regulatory activity encoded within individual 3’-UTRs.

## Results

### Distinct pathways regulate maternal mRNA decay

Fertilization triggers a remodeling of the transcriptome required for the first steps of embryonic development. Maternal and zygotic post-transcriptional pathways regulate the dynamics of mRNA stability during the Maternal-to-Zygotic transition (MZT) (Yartseva et al, 2015). The post-transcriptional regulatory code of proteins and RNA elements controlling mRNA stability during MZT remain poorly understood. To understand this regulation, we analyzed transcript levels during the first 8 hours post-fertilization (hpf), at 30-60 minute intervals in wild type zebrafish embryos using mRNA-seq. We identified mRNAs whose decay was dependent on the zygotic transcription or specifically on miR-430. These transcripts were stabilized when zygotic transcription was inhibited with the RNA polII inhibitor, *α*-amanitin (Lindell et al, 1970; Kane et al, 1996), or when miR-430 was inhibited using an antisense tiny-LNA complementary to miR-430 (LNA^430^) (Staton et al, 2013). To normalize RNA expression across stages we used exogenous yeast spike-in RNA, allowing us to quantify global changes in mRNA levels (Figure 1A). Comparing early (2 hpf) to late (6 hpf) developmental stages, we defined the main regulatory program for 5847 mRNAs undergoing decay: 3909 were regulated by the zygotic program, of which 616 were primarily dependent on miR-430 (11% of total unstable); and 1938 were primarily regulated by the maternal program (33%) (Figure 1B). *In situ* hybridization analysis of endogenous transcripts selected from each target set displays stabilization patterns in the absence of zygotic transcription (*α*-amanitin) and/or miR-430 function (LNA^430^) that were consistent with behaviors observed in the global RNA-seq analyses (Figure 1C).

**Figure 1.**
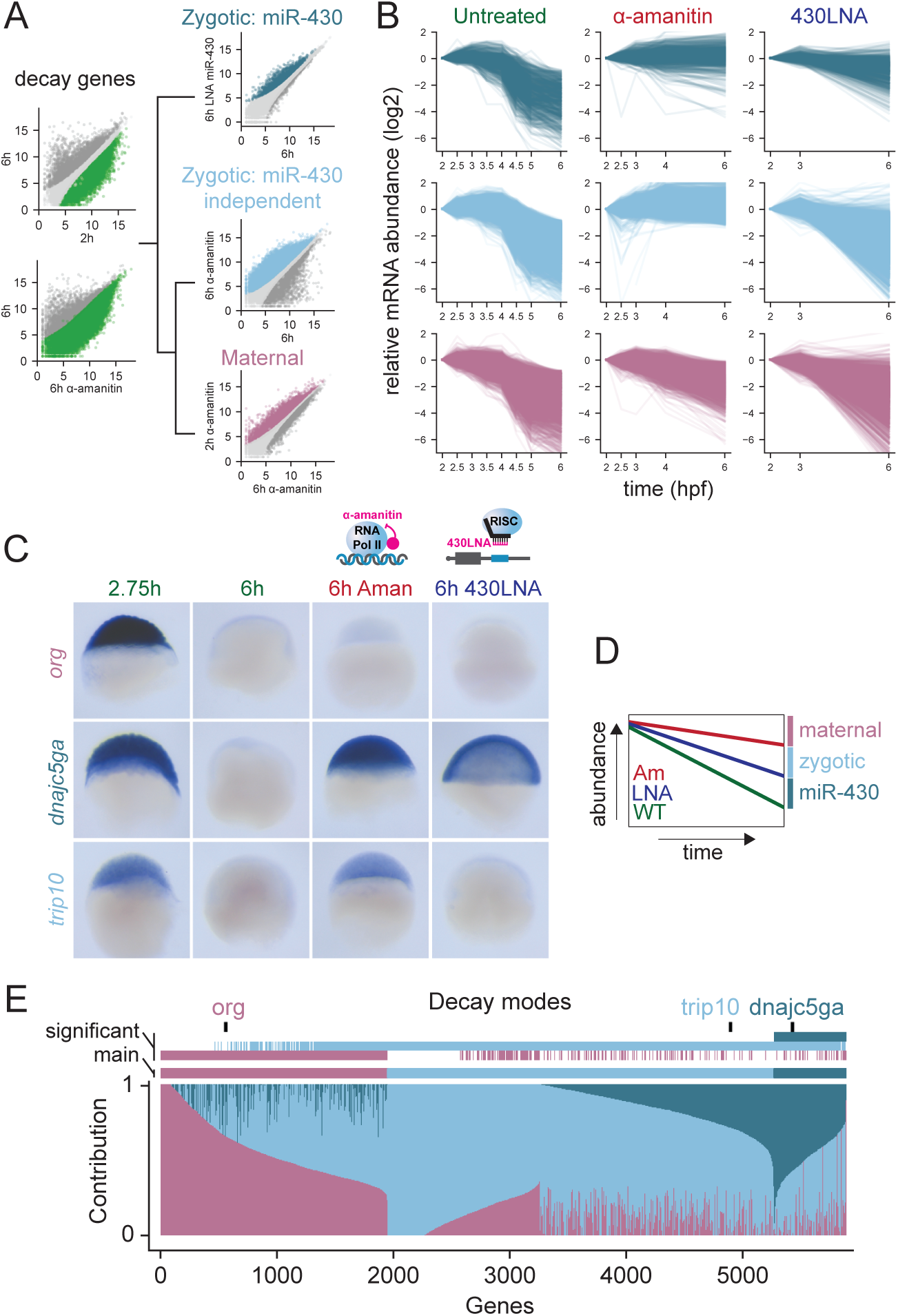
Distinct pathways regulate maternal mRNA decay. (A) Hierarchical protocol followed to separate genes into their predominant mode of decay. Among decaying genes determined as stable early (2 hpf) or higher in absence of zygotic transcription (*α*-amanitin treatment), miR-430 decaying genes were first characterized using LNA^430^ treatment. Remaining genes, as indicated by the black tree, were split into maternal and zygotic decaying genes based on their dependence on zygotic transcription. (B) Biplots representing mRNA expression levels over developmental time separated by decay pathways. Timecourse is shown in WT, *α*-amanitin and LNA^430^ conditions. (C) *In situ* hybridization illustrating three genes from the maternal, zygotic and miR-430 dependent decay modes. (D) Relative contribution of the three pathways to RNA decay. Fold-changes calculated in (A) were normalized taking into account *α*-amanitin treatment is also blocking miR-430 transcription. For example, if a transcript’s decay profile was not affected by blocking miR-430 or zygotic transcription, then that transcript’s degradation was wholly attributed to maternal decay program. In contrast, decay that was prevented by blocking miR-430 or zygotic transcription was attributed to miR-430 or zygotic decay pathways, respectively.

Although we were able to identify the predominant mode of decay for each unstable mRNA, the stability of any individual mRNA is likely dictated by multiple regulatory mechanisms operating within the embryo. This was reflected in the overlap between regulatory modes identified for numerous mRNAs. To dissect potential combinatorial regulation, we calculated the degree of stabilization conferred on individual mRNAs by each experimental treatment (loss of miR-430 regulation, or zygotic transcription; Figure 1D; Supp. Table 1). We find that the majority of unstable transcripts displayed decay behaviors indicative of combinatorial regulation by multiple decay pathways (Figure 1E). For example, we observe that up to 3688 mRNAs (63%; >0 contribution) are only partially stabilized after blocking miR-430 function. Together, these results define three regulatory modes of maternal mRNA turnover and their relative contributions to shaping post-transcriptional regulation across the zebrafish transcriptome after fertilization.

### RESA identifies regulatory RNA elements

We hypothesized that the combinatorial regulation observed for maternal transcripts might be encoded in discrete elements within the mRNA. To identify these elements, we used RESA (Yartseva et al, 2017; Figure 2A), which assesses the ability of sequences to regulate mRNA stability *in vivo* when placed in the 3’-UTRs of a reporter mRNA library. We generated two reporter libraries in parallel, one composed of random fragments spanning the entire embryonic transcriptome (~30 nt length; transcriptome library) and a high-density library enriched for 3’-UTRs from 434 genes regulated during the MZT (~100 nt length; targeted library). To identify regions that mediate differential mRNA stability, we quantified the abundance of each reporter within the RESA library before and after zygotic transcription in wild type (2 hpf, 6 hpf), *α*-amanitin or LNA^430^ injected zebrafish embryos. Depletion or enrichment of sequences over developmental time revealed 1404 destabilizing and 295 stabilizing regions respectively across 3456 genes. Destabilizing regions were modulated by either the maternal (593), the zygotic (184) or the miR-430 (627) programs of mRNA decay (Supp. Figure 3A). To test the regulatory activity of these sequences, we validated two reporter mRNAs containing 3’-UTR sequences identified by RESA using qRT-PCR (Figure 2B, C). Each reporter was destabilized in wild type embryos, and this effect was blocked when inhibiting zygotic transcription, consistent with the specific regulation of these regions by the zygotic program. We observed that the mean destabilization across miR-430 target sites corresponded to the predicted microRNA target site strength (8mer>7mer>6mer) while inserts containing reverse complement miR-430 target sites were not depleted (Figure 2D), demonstrating that RESA can accurately quantify regulatory strength across target sites.

**Figure 2.**
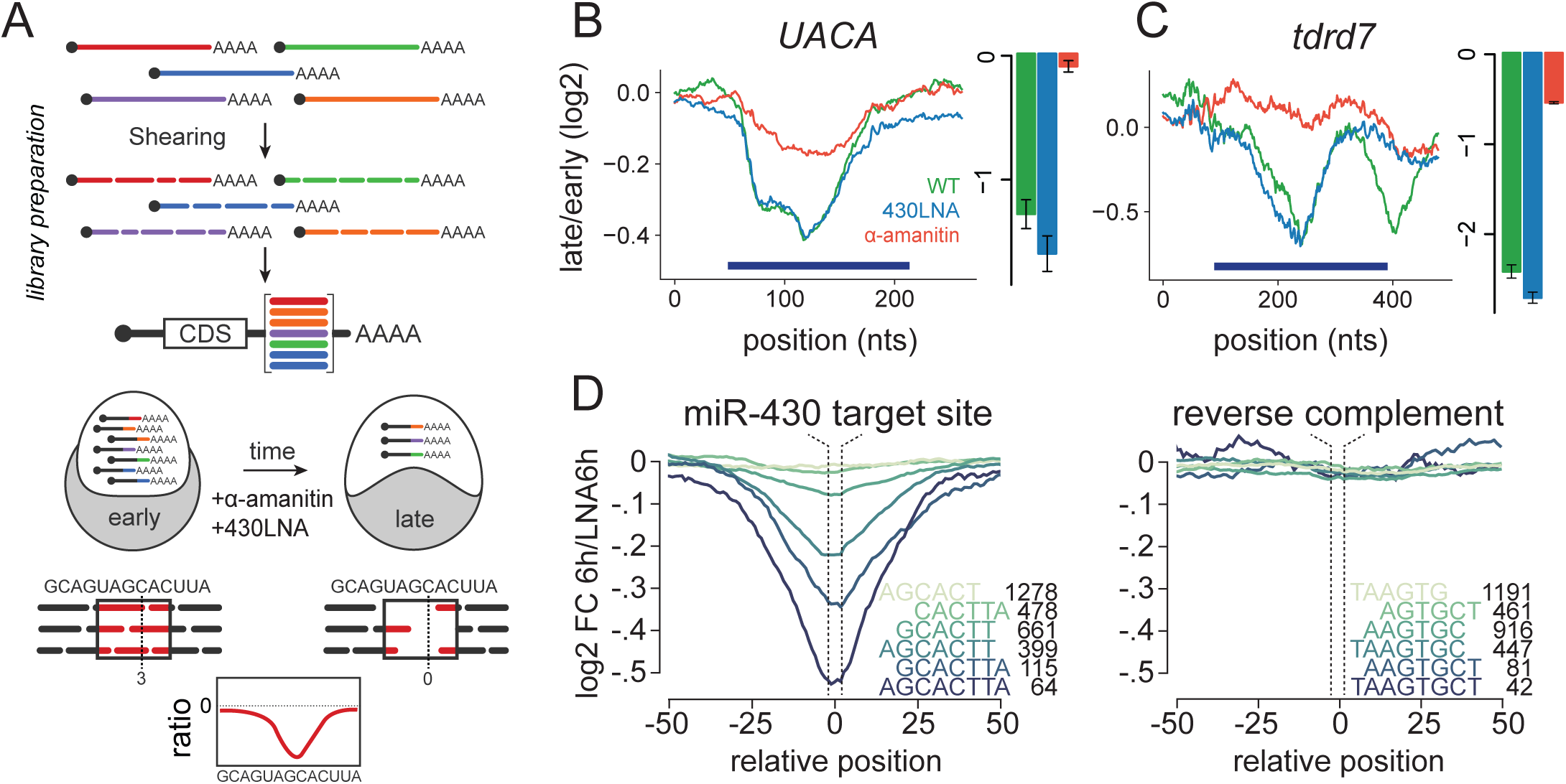
RESA identifies regulatory RNA elements. (A) Schematic of RESA method. Sequence fragments derived from endogenous transcripts are placed within the 3’-UTR context of reporter mRNAs. Regulatory activity is inferred from relative depletion or enrichment post-injection. Treatment with *α*-amanitin or 430-LNA delineates sequences under zygotic and/or miR-430 regulatory pathways, respectively. (B-C) RESA identifies sequence regions within *UACA* and *tdrd7* transcripts that are responsive to zygotic regulatory mode. Bar graphs display independent validation by RT-PCR with reporters containing sequence inserts spanning regulated regions (dark blue bars). Late/early fold change in untreated (green), 430-LNA-treated (blue), *α*-amanitin treated (red). (D) Mean destabilization across all loci (min coverage of 5 counts per million at 2 hpf) centered on miR-430 target site variant, with number of represented loci indicated for each variant. Right panel shows mean destabilization for loci possessing the reverse complement for each miR-430 target sequence.

**Figure 3.**
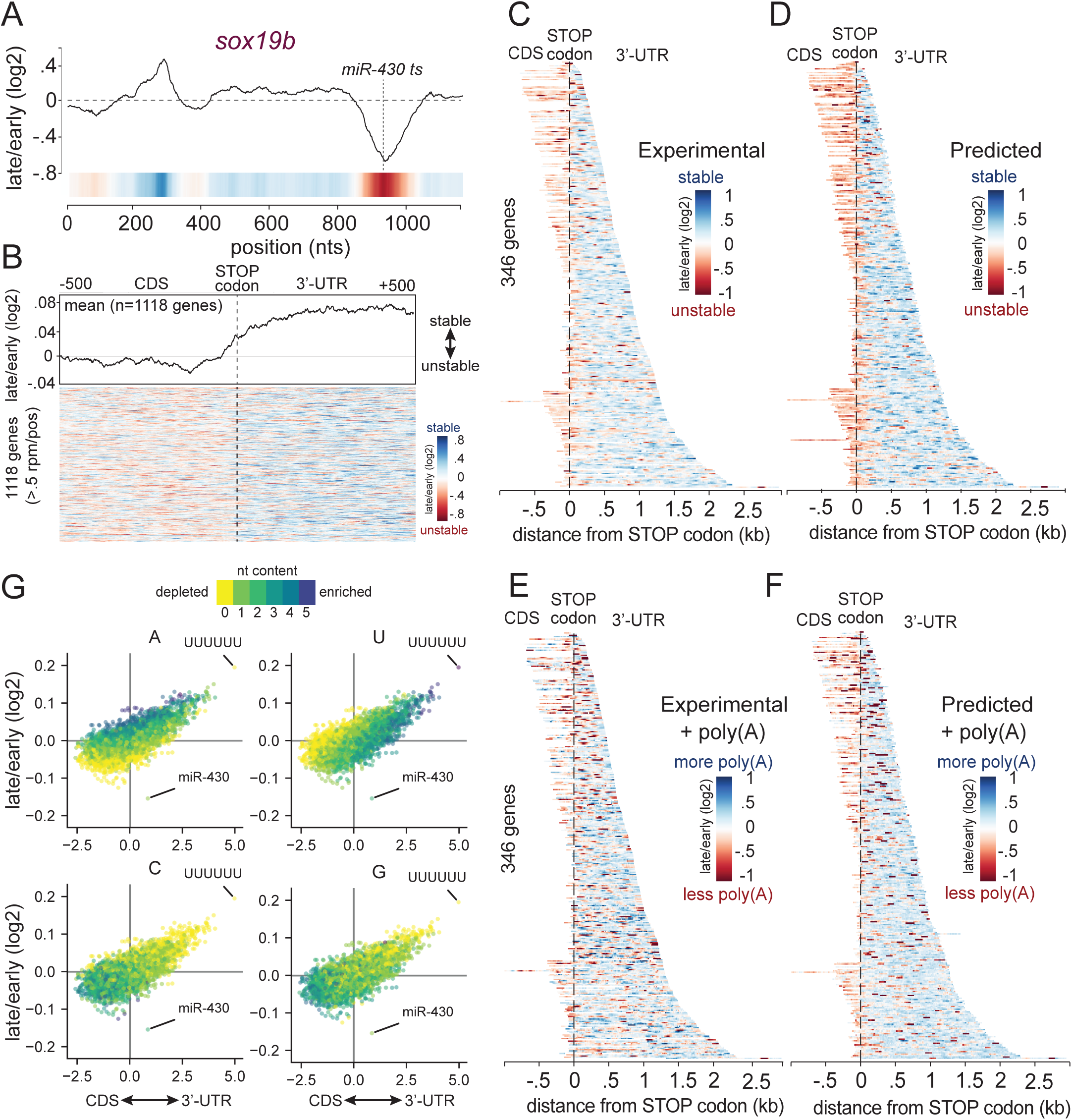
Nucleotide composition underlying CDS and 3’-UTR sequences impacts mRNA stability. (A) RESA coverage ratio for the *sox19b* locus. The RESA-verified miR-430 target site marked with dashed lines. (B) Heatmap illustrating transcriptomic RESA coverage ratios of 1118 genes above 0.5 CPM per position within 1 kb window centered on STOP codon. Averaged signal is represented in the top plot illustrating the higher stability of 3’-UTR derived sequences. (C-F) Heatmap illustrating the coverage ratio over the targeted RESA total RNA (C, D) and poly(A) selected libraries (E,F) respectively. For each of these two libraries, (C, E) shows the experimental coverage ratio (>log2(10) CPM), while (D, F) shows the predicted coverage ratio according to the Random forest model trained on each of these two libraries separately. (G) Biplots comparing all hexamers relative enrichment within CDS and 3’-UTR to their average coverage ratio between 2 hpf and 6 hpf. From left to right, biplot is colored with number of U, A, C and G within each hexamer.

### Nucleotide composition underlying CDS and 3’-UTR sequences impacts mRNA stability

RESA identified hundreds of stabilizing and destabilizing regions in mRNAs. We observed that stabilizing and destabilizing sequences frequently reside on the same transcript. For example, the 3’-UTR of *sox19b* harbors two regulatory sequence elements, one stabilizing and one destabilizing (Figure 3A), which could potentially act antagonistically in affecting mRNA half-life, or could be temporally regulated.

To understand the features that characterize regions with stabilizing and destabilizing effects on the mRNA, we first compared the regulatory strength of each sequence with respect to its position within the endogenous transcript. Sequences derived from 3’-UTR regions tended overall to have stabilizing effects (with exceptions notably miR-430 target sites), whereas CDS-derived sequences were significantly more destabilizing when placed in a 3’-UTR context (transcriptome-wide, Figure 3B, G; Pearson r=0.70; targeted RESA libraries, Figure 3C, G; Pearson r=0.77). Across all the sequences, sites complementary to miR-430 confer the strongest destabilizing effect. Because RESA measures the activity of sequence elements when placed within a uniform 3’-UTR location, these results suggest that intrinsic features of these regions account for the differential effect on mRNA stability.

During early embryogenesis, deadenylation and decay are uncoupled (Voeltz et al, 1998), allowing us to distinguish between elements that predominantly cause deadenylation or decay. We compared total RNA versus poly(A)-selected targeted RESA libraries, extracted after injection. Relative poly(A) tail length is inferred from capture efficiency (i.e depletion of reads from poly(A) selection over time relative to total reads indicates deadenylation). We observed that CDS and 3’-UTR derived sequences overall remained destabilizing or stabilizing respectively (Figure 3E). Within the strongest destabilizing elements in the 3’-UTRs, microRNAs confer both deadenylation and decay of the across both libraries (total and poly(A) selected). In contrast, other regions were depleted in poly(A) selected but not in the total mRNA library, suggesting that specific sequences have differential effect on mRNA deadenylation and stability.

To identify specific sequence features enriched within the regulatory regions identified in RESA, we analyzed their nucleotide composition and their position bias in the mRNA. We compared the mean regulatory activity of all possible hexamers to their prevalence within 3’-UTR versus CDS regions (Figure 3G). Uridine or adenine-rich hexamers enriched in the 3’-UTRs (correlation between U-content and 3’-UTR hexamer enrichment: r=0.59; A: r=0.15; Supp. Figure 1A) were stabilizing *in vivo* (correlation between U-content and hexamer RESA signal U: r=0.33; A: r=0.60; Supp. Figure 1B), with polyuridine sequences (UUUUUU) displaying the strongest stabilization. In contrast, cytosine or guanine-rich hexamers are predominantly found within CDS regions (anti-correlation between C-content and 3’-UTR hexamer enrichment: r=-0.38; G: r=-0.28; Supp. Figure 1A) and tended to be destabilizing (anti-correlation between C-content and hexamer RESA signal C: r=-0.35; G: r=-0.28; Supp. Figure 1B). These results were confirmed using the targeted RESA library (Supp. Figure 1C-E). They suggest that nucleotide composition bias underlying CDS and 3’-UTR sequences is associated with differential RNA stability.

### Identifying destabilizing and stabilizing regulatory motifs

To identify short-linear motifs enriched in co-regulated sequences, we used FIRE (Finding Informative Regulatory Elements) (Elemento et al, 2007; Oikonomou et al, 2014). This method analyzes all possible 7-mers, to then optimize these seeds into more general motif representations by maximizing mutual information (Elemento et al, 2007; Oikonomou et al, 2014). We identified motifs associated with destabilization and stabilization in the three decay modes (Figure 4A; Supp. Figure 2; confidence cutoff Z-score > 20 and > 10 for the transcriptome and targeted libraries respectively, see Methods). We find that miR-430 seed target sequences were specifically enriched in unstable sequences that were stabilized by the loss of miR-430 function. Independent of miR-430, the motifs most strongly associated with unstable sequences were CCUC and CUGC (Z-score 144.1 and 77.8). To validate these findings independently from RESA, we analyzed expression of a reporter mRNA containing multiple copies of the CCUC motif derived from the gene *zc3h18*. We observed decreased stability for CCUC motifs compared to a reporter where those motifs were mutated to CGUC, resulting in lower protein expression of a GFP reporter mRNA compared to a dsRed control (Figure 4B-D). These results suggest that CCUC motifs are responsive sequence-specific decay elements in the early zebrafish embryo. FIRE identified poly-U motifs such as UUUNUUU, UUUUANA or AANUAUU over-represented in stable regions with poly-U displaying the strongest activity (Supp. Figure 2). Stabilization at multiple poly-U sites was consistently observed within individual transcripts (Figure 4E). In contrast, poly(A) enriched samples revealed that deadenylated sequences were enriched in miR-430 sites and U[CA]UUUAUU, an AU-rich element (ARE) known to be involved in regulating poly(A) tail length (Audic et al, 1997; Voeltz et al, 1998; Yartseva et al, 2017; Rabani et al, 2017) (Figure 4F; Supp. Figure 2). To validate this motif, we analyzed the polyA tail length of a reporter mRNA containing tandem copies of UAUUUAUU derived from the gene *fam116b.* This analysis revealed that poly(A) shortening was dependent on activation of the zygotic genome, and abolished in *α*-amanitin treated embryos (Figure 4G).

**Figure 4.**
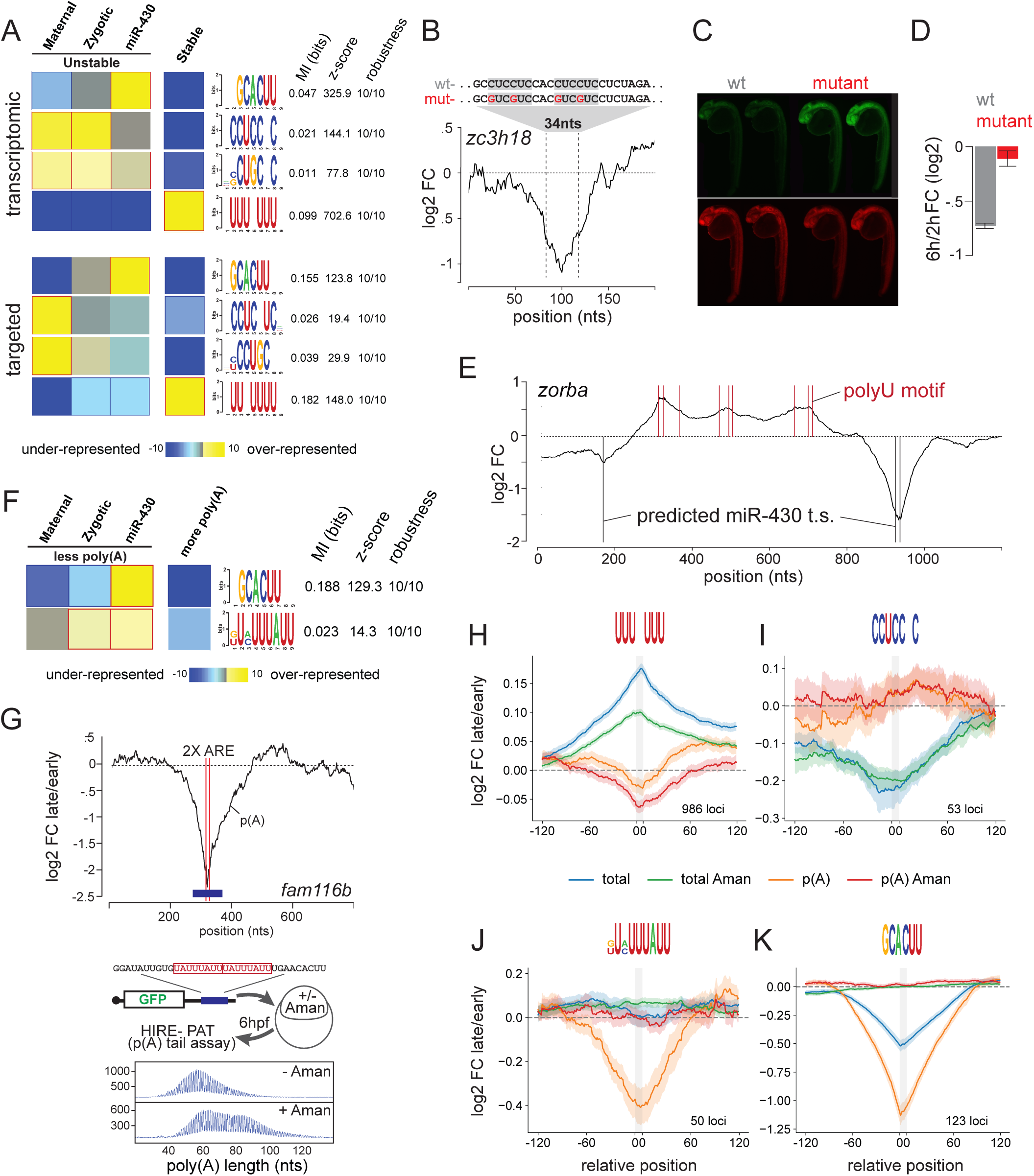
Identifying destabilizing and stabilizing regulatory motifs. (A) FIRE top selected motifs. In heatmap, overrepresentation (yellow) and underrepresentation (blue) patterns are shown for each discovered motif in the corresponding category. Also shown are the mutual information values, Z-scores associated with a randomization-based statistical test and robustness scores from a 3-fold jackknifing test. (B) RESA coverage ratio for the *zc3h18* locus zoom on peak containing multiple copies of the CCUC motif. (C) Zebrafish 24h embryos injected with WT (CCUC motifs) or mutant (CGUC motifs) reporters. (D) Bar graphs displaying independent validation by RT-PCR. (E) RESA coverage ratio for the *zorba* locus. Poly-U and miR-430 target site are marked with read and black bars respectively. (F) Same as (A) for poly(A) selected RESA library. (G) RESA with poly(A) selection coverage ratio plot showing the two ARE sites (red lines) within the *fam116b* locus (top). HIRE-PAT assay measuring impact of *fam116b* ARE sites on poly(A) tail length reporter. Shortening of poly(A) tail length is zygotic transcription dependent as shown by longer poly(A) tail after *α*-amanitin treatment (bottom). (H-K) Motif-centered metaplots for poly-U (H), CCUCCNC (I), ARE (J) and miR-430 (K) motifs. Targeted RESA libraries coverage ratio were averaged over windows centered on RBP motif (RESA minimum coverage >0.01 CPM). Motif is represented with grey bar representing standard error of the mean (S.E.M.) of RESA is shown by shaded outlines.

To measure the regulatory activity of each identified motif, we analyzed the average decay across all the regions containing that motif in the RESA assay before (2hpf) and after MZT (6-8hpf). To gain insights into the regulatory mechanism for each motif we performed two different assays: i) we compared their effects in the total and the poly(A) selected library to distinguish effects on stability and deadenylation, and ii) we analyzed their dependency on the zygotic mode of decay by determining whether their regulatory activity was blocked in the absence of zygotic transcription (*α*-amanitin treated samples). Poly-U motif showed a stabilizing effect (mean stabilization of 0.17 log2 fold-change at 2 vs. 6 hpf; Figure 4H), while CCUCCNC had a destabilizing effect (mean destabilization -0.22 logFC 6/2hpf; Figure 4I). Regulation by these elements was mainly controlled through the maternal program on the total mRNA as it was still observed in *α*-amanitin treated samples, but had less effect on the poly(A) libraries. Analysis of lower Z-score FIRE motifs CCUGC and UUAUU, and Pumilio (Gamberi et al, 2002; Gerber et al, 2006) motifs revealed weak regulation acting on total and poly(A) libraries (Supp. Figure 3B-D). While these elements could have intrinsically weaker effects, this could suggest combined regulation of multiple motifs. As a control, meta-analysis over 123 miR-430 seeds (Figure 4K) showed a strong zygotic-dependent regulatory effect on stability (-0.5 logFC 6/2hpf) and deadenylation (-1.15 logFC 8/2hpf), with an effect that was > 2.5 times stronger than poly-U and CCUCCNC motifs. Consistent with the reporter analysis, meta-analysis of the U[CA]UUUAU ARE motif over 50 loci reveals that this motif causes zygotic-dependent deadenylation (-0.4 logFC 8/2hpf) with a robust depletion of polyadenylated fragments, without a significant effect on the total mRNA abundance within the time frame analyzed (Figure 4J). Relative regulatory activity of these elements revealed that within this context, miR-430 provides stronger regulation than ARE motifs UAUUUAUU, with a rapid coupling of deadenylation and decay that is not observed in AREs. Together, these results identify sequence motifs that regulate reporter mRNA stability and deadenylation.

### Mapping RNA-interacting factors in the embryo

To identify the *trans*-factors binding the regulatory sequences identified by RESA and FIRE, we adapted the interactome capture technique to zebrafish embryos (Castello et al, 2012; Baltz et al, 2012; Kwon et al, 2013; Wessels et al, 2016; Sysoev et al, 2016; Despic et al, 2017). This method uses UV crosslinking of protein-mRNA interactions and poly(A) purification followed by mass spectrometry to identify the proteins bound to mRNA (Figure 5A, Supp. Figure 4). We analyzed the interactome across three independent biological replicates, with or without UV crosslinking. We queried the interactome at 4 hpf, a timepoint that precedes the widespread changes in maternal mRNA stability characterized here, when RBPs are already engaged in RNA-binding (Bazzini et al, 2012). Using label-free quantitative mass spectrometry, we identified 160 proteins with ≥ 2 peptides with at least one of them unique. 112 of the 160 identified proteins were significantly enriched in the UV crosslinked sample compared to controls (Figure 5B). From this core set of 112 proteins, 90 had been also identified in previous mRNA interactomes (Castello et al, 2012; Baltz et al, 2012; Kwon et al, 2013; Liao et al, 2016; Despic et al, 2017) (Supp. Table 2). We observed a significant enrichment of RNA binding proteins as 67.8% of the identified proteins had annotated RNA-binding domains compared to 8.2% of proteins detected in the input. Our analysis reveals an enrichment for Gene Ontology terms associated to RBPs (Figure 5C), and a selective enrichment for *bona fide* RNA-protein interactions (Supp. Figure 5). These included RBPs involved in RNA processing and splicing such as *xrn2*, *srsf2* or *celf1*, in proteins controlling mRNA translation such as *eif4a*, *eif2a* and *eif4enif1*; and RNA stability such as *pumilio*, *piwi*, *hnRNPd*, *khsrp* and *staufen* among others. To validate their ability to bind RNA we immunoprecipitated six of the identified proteins after UV crosslinking, and radio labeled the bound RNA for detection (Figure 5D, E). While all the proteins interacted with RNA, a subset of these interactions was also dependent on the activation of zygotic transcription (*ythdf1*, *khsrp*, *khdrbs1*), since the levels of RNA pulled down were reduced in *α*-amanitin treated embryos. Taken together, the zebrafish interactome identify a large set of RBPs that participate in wide range of RNA processing pathways during the MZT, providing an entry point to identify the effector proteins that regulate mRNA stability and decay.

**Figure 5.**
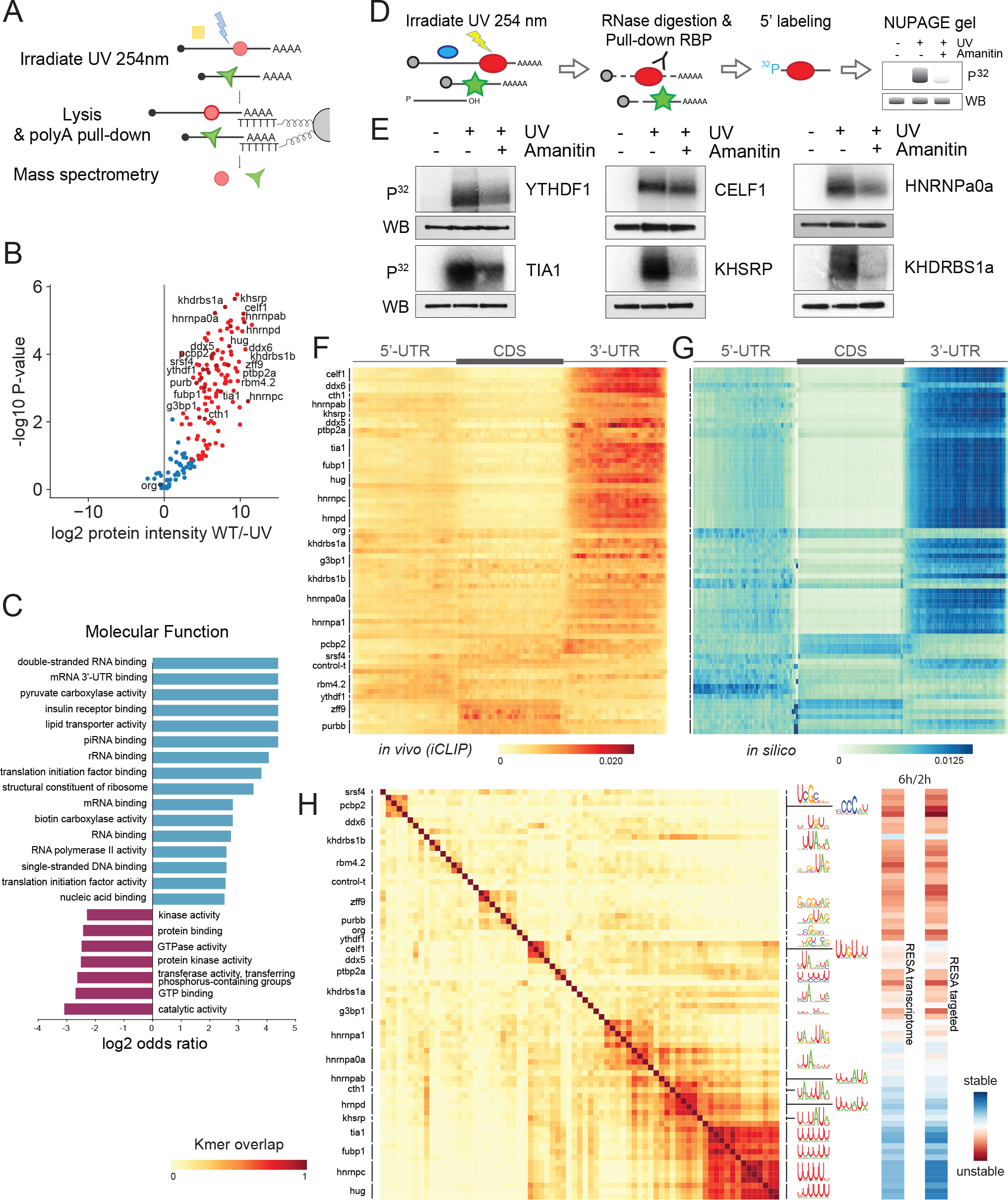
Mapping RNA-interacting factors in the embryo and identifying putative regulatory factors driving mRNA stability. (A) Diagram summarizing the interactome capture protocol. After UV-crosslinking and poly(A) mRNA pull-down, RBPs are identified using mass spectrometry. (B) Volcano plot showing the RBPs significantly enriched over background by interactome capture. (C) GO term enrichment analysis characterizing the molecular functions of the captured proteins. (D) Cartoon depicting the rationale behind a label-transfer experiment to validate RNA-protein interactions. P^32^ autoradiograph indicates the amount of RNA while FLAG Western blot indicates RBP levels. (E) Validation of the RNA-binding activity during zebrafish development of representative RPBs identified in the interactome capture. (F) Heatmap representing iCLIP metaplots of RBP-binding within protein-coding transcripts. Metaplots averaged over each RBP were clustered to group similar binding profiles. (G) Heatmap representing *in silico* binding profiles obtained by scanning for RBP binding motifs within protein-coding transcripts. Motifs are represented in (H). (H) Heatmap illustrating the overlap among RBP binding motifs. Motifs were characterized using the top most 6-mers bound normalized by iCLIP control (left). RESA averaged coverage ratio for each motif (right).

### Identifying putative regulatory factors driving mRNA stability

RESA identified strong regulatory differences between 3’-UTR and CDS-derived sequences that we associated with different nucleotide composition. To identify potential effector proteins that mediate the regulatory activity observed in RESA, we performed iCLIP experiments on 24 RBPs identified in the zebrafish interactome and analyzed their target sequences (Supp. Table 3). We reasoned that having a common tag would allow us to compare the signal between different proteins to identify specific binding events for each protein and distinguish them from background common to all samples. Thus, we analyzed FLAG-tagged versions of each protein expressed from an injected mRNA. To ascertain that this approach captures *bone fide* binding sites, we compared iCLIP signal of one candidate, *khsrp*, obtained with the FLAG-tagged protein version or using an antibody to pull down the endogenous protein. We saw a similar enrichment of both the FLAG-protein and the endogenous *khsrp* in the 3’-UTR of endogenous mRNAs and a common motif enriched in both analyses among the iCLIP target sequences (motif UUUAU[AU]) (Supp. Figure 6).

**Figure 6.**
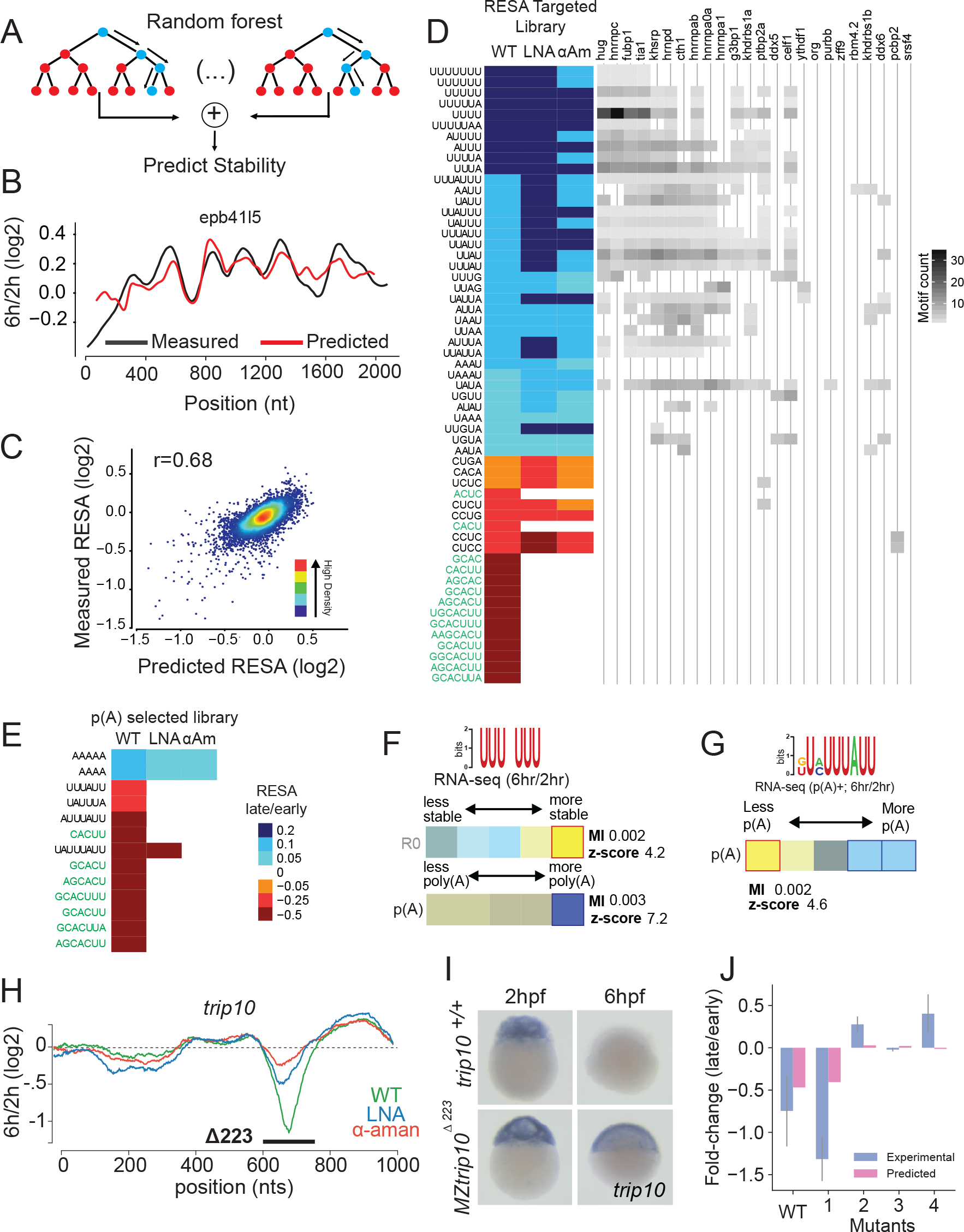
Modeling the effect of sequence on mRNA stability in vivo. (A) Scheme of the procedure for building the random forest model on RESA profiles. Data generated from window-sliding across the 3’-UTRs are used to train a Random forest model. (B) Example of predicted (red) and RESA experimental (black) profiles for *epb41l5* gene. (C) Model performance per window using 5-fold cross validation; Model achieved 0.68 Pearson correlation between predicted stability (x-axis) and measured RESA stability (y-axis). (D) (left) Top selected motifs according to the random forest model trained on RESA targeted library. Each column represents WT, LNA and *α*-amanitin treatment respectively. Color intensity represents the RESA fold-change difference between windows that contain or not each motif. Blue represent stabilizing motifs and red represent motifs associated with decay (all motifs have *P*-values below 4.6e-56 (Mann-Whitney U-test followed by Bonferroni multiple testing adjustment). (right) Heatmap representing motif enrichment in the top 50 hexamers enriched in iCLIP experiments. (E) Same as (D) with random forest model trained on the poly(A) selected RESA library. (F, G) poly-U (F) and ARE (G) motifs enrichment between 2 hpf and 6 hpf within total and/or poly(A) selected RNA-seq. In heatmap, overrepresentation (orange/yellow) and underrepresentation (blue) patterns are shown. Also shown are the mutual information values, Z-scores associated with a randomization-based statistical test and robustness scores from a 3-fold jackknifing test. (H) RESA profile from the *trip10* locus. Genetic deletion of a sequence spanning regulated region (black line) results in stabilization of the trip10 transcript as assessed by *in situ* hybridization (I). (J) Random forest model validation. Barplot comparing experimental and predicted average stability of *trip10* decay peak and four mutated sites (Spearman r=0.60).

We characterized the binding pattern and the sequence motif preferentially bound by each RNA binding protein across replicates. Cumulative count of the iCLIP reads within the 5’-, 3’-UTRs and CDS, revealed that the majority of RBPs displayed strong occupancy within the 3’-UTR (Figure 5F). Within this class, we observed variable accumulation of reads in different regions of the 3’-UTR. Among these, *celf1*, *hug*, and multiple hnRNPs displayed preferential occupancy towards the distal end of the 3’-UTR. In contrast, *pcbp2* binding was more frequently observed directly proximal and downstream to the annotated stop codon. iCLIP reads from *zff9* and *purbb* were distributed throughout the CDS. In particular, *khsrp* was preferentially enriched across the 3’-UTRs in the transcriptome and was preferentially excluded from coding sequences and 5’-UTRs. Next, we assessed the RNA sequence specificity of each RBP by searching for the top 10 enriched hexamers bound by each RBP compared to a negative control lacking a FLAG protein (Figure 5H). Proteins that had a motif previously identified *in vitro* (Ray et al, 2013), showed a similar sequence specificity, and a strong correspondence between *in vitro* and *in vivo* identified motifs (Supp. Figure 8). The iCLIP binding pattern for each RBP resembled the distribution of the top identified motifs (Figure 5G), suggesting that for most proteins, the presence of the binding motifs explains the binding distributions observed *in vivo*. However, we observed (i) higher density of *in silico* binding in the 5’-UTR than observed *in vivo* for several RBPs, and (ii) higher density of *in silico* binding to the CDS for *pcbp2* despite similar *in vivo* and *in vitro* profiles for *pcbp2* within the 3’-UTR. These differences raise the possibility that additional factors, such as the ribosome, might contribute to the observed occupancy profiles *in vivo* (Supp. Figure 7), as it has been suggested for UPF1 (Zünd et al, 2013).

**Figure 7.**
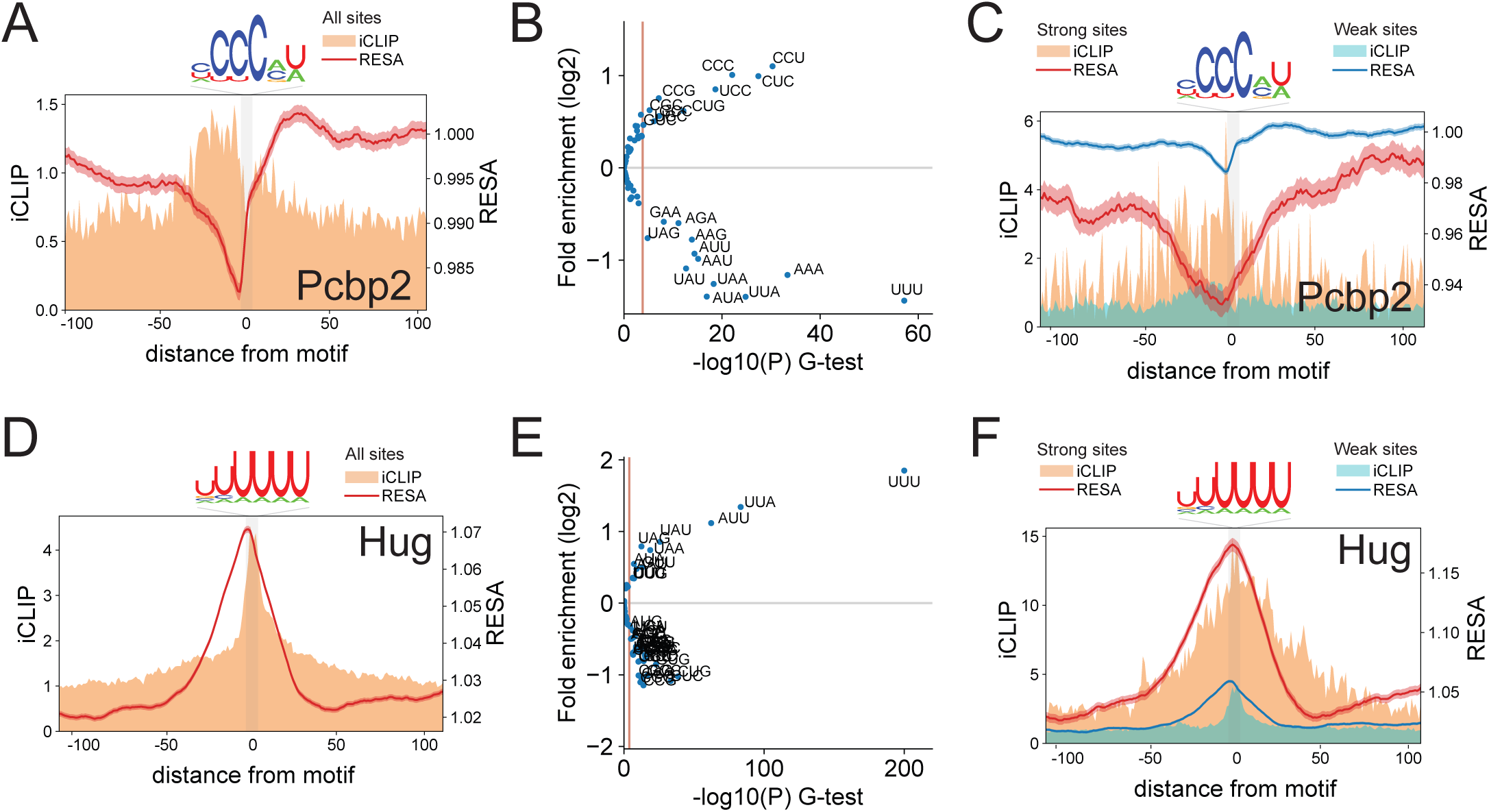
Antagonistic effects of different RBP binding motifs. (A, D) Motif-centered metaplots for *pcbp2* (A) and *hug* (D). RESA coverage ratio and iCLIP signal were averaged over windows centered on RBP motif (RESA minimum coverage >0.05 CPM). Motif is represented with grey bar. S.E.M. of RESA is shown by red shaded outline. (B, E) Sequence context of RBPs potential target sites. Volcano plot representing 3-mer enrichment 20 nt upstream and downstream motif between the top 10% most destabilizing for *pcbp2* (B) and most stabilizing for *hug* (E) and the bottom 10%. *P*-values were calculated using G-test. Red line indicates 1% significance cutoff after Bonferroni multiple test correction. (C, F) Motif-centered metaplots for *pcbp2* (C) and *hug* (F) comparing potential target sites with favorable context (red) or not (blue). Favorable context defined as top 10% of sites most enriched with significantly enriched 3-mers and least enriched in significantly depleted 3-mers as shown on panels B and E. Scale on y-axis was adjusted showing higher regulation and wider iCLIP binding signal of sites with favorable context.

We found that several of the RBP binding motifs possess regulatory activity as measured by RESA (Figure 5H; Figure 6D; right panels). For example, *hug*, *fubp1*, *hnrnpc* and *tia1* were preferentially bound to poly-U associated with stabilizing motifs. In contrast, RBPs such as *pcbp2*, *ptbp2a* or *zff9* were preferentially bound to destabilizing motifs. Taken together, these data link the regulatory activity of specific RNA motifs to the recognition by specific RBP during zebrafish embryogenesis.

### Modeling the effect of sequence on mRNA stability in vivo

Our analysis identifies several motifs that are enriched within regulated sequences in the mRNA. To determine whether sequence information can be used to model reporter mRNA regulation, we used machine learning and developed a random forest model (Breiman, 2001). We reasoned that this model would allow us to capture the association between motif frequencies and their corresponding regulatory activities along RNA sequences (Figure 6A). We analyzed the frequency of kmers (1-8 nt) in 100 nt sliding windows across the RESA library. To build the model, we retained 387 kmers out of 87,380 with absolute Spearman’s rank correlation coefficient above 0.1 between kmer frequency and fold-change. Next, we used 5-fold cross validation to train and assess the performance of the random forest model. This model achieved a 0.68 Pearson correlation between the predicted change in mRNA abundance for each window, and the corresponding change measured by RESA (Figure 6B, C). This analysis selected 57 motifs with a dominant effect on stability (Figure 6D; left panel, Supp. Figure 10A).

To further analyze the importance of the motifs selected by the random forest model, we compared the differential stability between windows that contain or lack each motif. This analysis revealed that the top 25% stabilizing windows were significantly enriched in poly-U motifs, as well as UUAG and UGUA (*P* < 1e-146, 8e-21 and 9e-20 respectively, Mann-Whitney U-test with Bonferroni multiple testing adjustment), echoing the trends observed above. On the other hand, the top 25% destabilizing windows were significantly enriched in miR-430 complementary sites and CCUC motifs (*P* < 2e-96 and 6e-95 respectively, Mann-Whitney U-test with Bonferroni multiple testing adjustment) (Supp. Figure 9B-C). By intersecting these kmers with the sequences identified in the iCLIP experiment, we identified potential RBPs that could act as *trans*-factors to regulate bound mRNAs (Figure 6D; right panel) and revealed UUAG was significantly enriched among the top 50 kmers bound by *hnrnpa1* (*P* < 2e-73, χ^2^-test followed by Bonferroni multiple testing adjustment) and UUUU was significantly enriched in kmers bound by *hug* and *hnrnpc* (*P* < 1e-174 and 2e-92 respectively). On the other hand, CCUC was significantly enriched in kmers bound by *pcpb2* (*P* < 2e-23) and CUCU was significantly enriched in kmers bound by *ptbp2a* (*P* < 1e-30). Together these analyses provided a model that predicts the regulatory information encoded in the 3’-UTR sequence to regulate reporter mRNA stability *in vivo*.

mRNAs can be regulated by maternally or zygotically encoded programs. To model the different modes of mRNA decay, we built additional random forest models in the absence of zygotic regulation (*α*-amanitin) or miR-430 regulation (LNA^430^). We observed a strong correlation between the predicted and the observed regulation in RESA for both models (*α*-amanitin, r=0.70, Supp. Figure 10B; LNA^430^ r=0.74, Supp. Figure 10C). Consistent with the FIRE analysis, CCUC and poly-U motifs were identified as regulatory sequences independent of zygotic transcription as part of the maternal model, while miR-430 was identified as the main element regulating mRNA stability of the zygotic mode (Figure 6D). In contrast, models built using poly(A) selected libraries were less accurate (wild type, r=0.53; *α*-amanitin, r=0.33; LNA^430^, r=0.31, Figure 6E, Supp. Figure 11) and revealed that UAUUUAUU (AREs) motifs were the strongest zygotic-dependent regulator of the poly(A) tail after miR-430. The lower accuracy of the model when the zygotic mode is blocked suggests that 3’-UTR sequence mediated regulation of the polyadenylation status is mainly regulated by zygotic-dependent mode. Based on these results, we conclude that our current models capture the importance of CCUC and poly-U to maternally regulate RNA stability and define miR-430 and UAUUUAUU (AREs) to zygotically regulate the polyadenylation status of the mRNA.

Finally, we extended the training data to library sequences originating from coding-sequences (Figure 3D, F; total and poly(A) selected libraries respectively). We observed these new models predicted the global CDS/3’-UTR pattern and individual regulatory profiles we observed *in vivo*. These results suggest that our model was able to capture the intrinsic and position-biased features of these regions driving mRNA stability and poly(A) tail length.

### Predicting stability of endogenous mRNAs

Because the random forest captures the association between motif frequencies and stability, we applied it to predict differential stability of endogenous mRNAs. However, our model only achieved a 0.29 Pearson correlation between the predicted and measured mRNA fold changes (Supp. Figure 9D), suggesting that a large fraction of the regulation is not captured by the model developed on the 3’-UTR reporter mRNAs. Despite these limitations, regulation for 18% of the mRNAs was accurately predicted (fold-change within 10% of observed). These mRNAs were significantly enriched in motifs identified in RESA analysis (e.g. miR-430 and U-rich motifs, Mann-Whitney U-test with Bonferroni corrected P<0.05) (Supp. Figure 9E).

Among the RESA enriched motifs, several were associated with differential stability or deadenylation of endogenous mRNAs. Endogenous mRNAs containing poly-U motifs were specifically stabilized in total RNA libraries (Figure 6F). In contrast, we find that UAUUUAUU sequences (AREs) are enriched within endogenous mRNAs that are deadenylated, consistent with a role for ARE in poly(A) tail shortening (Figure 6G). To test the regulatory activity of endogenous sequences identified by RESA, and the motifs identified by the random forest model, we used CRISPR-Cas9 editing to mutate a potential destabilizing sequence element identified in the *trip10* gene. We observed that a 223 nt deletion in the endogenous gene, targeting the region regulated in RESA, caused stabilization of the mutant mRNA compared to wild type, demonstrating that RESA can identify *de novo* functional regulatory elements *in vivo* (Figure 6H, I). Sequence analysis of the regulatory region in *trip10* 3’-UTR revealed multiple AUUUA, AUUA and AAAUAAA that when disrupted stabilized the mRNA (Supp Figure 12B-C) and increased protein output (Supp Figure 12D). Our model predicted the reporter RNA levels with 0.60 Spearman correlation compared to levels measured experimentally (Figure 6J; Supp Figure 12A). For a large subset of endogenous mRNAs, additional regulatory modules in the CDS (Presnyak et al, 2015; Bazzini et al, 2016; Mishima et al, 2016) or sequence modification (Batista et al, 2014; Zhao et al, 2017; Ke et al, 2017) are likely to account for part of the regulatory interactions observed *in vivo*. Nonetheless, for miR-430 and AU-rich targets, a significant part of the regulation is encoded in the 3’-UTR sequence and could be accurately predicted using machine learning.

### Antagonistic effects of different RBP binding motifs

The regulatory effect of any particular motif can be expressed as the average activity across hundreds of loci with that motif. For example, loci containing the *pcbp2* binding motif displayed a mean destabilization of 0.985 across 10,305 loci (Figure 7A). However, for most motifs, we observed a broad spectrum of regulation and RBP binding, suggesting that the mere presence of a given motif is not the unique determinant of regulatory activity or RPB binding. We hypothesized that the sequence context for each target site might explain differential regulation (RESA) and/or binding (iCLIP). To assess this, we ranked each locus possessing a *pcbp2*, miR-430 or *hug* binding motif according to its RESA activity. An analysis of flanking sequences revealed significant enrichment of specific 3-mers when comparing the most regulated versus least regulated loci.

For *pcbp2*, CCU, CUC and CCC were significantly enriched within flanking sequences of the most destabilized sequences, which was associated with increased *pcpb2* binding as shown by cumulative iCLIP signal (Figure 7B, C). In contrast poly-U rich sequences were significantly depleted within these sites. Conversely, the context of *hug* binding sites was significantly enriched in UUU 3-mers within the most stabilized sequences, which were also among the most abundantly bound sequences by *hug* (Figure 7D-F, Supp. Figure 13A-C). miR-430 binding sites did not appear to have a specific sequence bias associated with stronger regulation, consistent with the strength of the regulation being modulated by the size of the seed alone (Supp. Figure 13D, E). Favorable sequence contexts for *pcbp2* and *hug* resemble the original binding site, and have opposing nucleotide preferences. These results suggest that U-rich and poly-C binding proteins might antagonize each other’s activity with binding and sequence context influencing the stability of the mRNA.

## Discussion

Post-transcriptional regulation plays a major role in shaping gene expression during cellular transitions, where mRNAs from the previous state need to undergo repression and decay. During the MZT, maternal and zygotic programs are integrated to regulate mRNA abundance. Here, we identified i) *cis*-regulatory elements, using RESA, ii) *trans*-factor RNA-binding proteins (RBP), using interactome capture, and iii) RBP target sequences using iCLIP. We integrated these regulatory sequences into a random forest prediction model, which largely recapitulated 3’-UTR mediated regulation as measured by RESA. These findings integrate important aspects of the post-transcriptional regulatory code shaping mRNA stability *in vivo*.

Our implementation of RESA allowed us to measure the relative strength of regulatory sequences *in vivo*. We find that sequences enriched in the 3’-UTR are mainly promoting mRNA stabilization, while sequences that were derived from the CDS are mostly promoting mRNA decay, an effect that was correlated with different nucleotide content. Among the regions in the 3’-UTR, miR-430 targets were the strongest deadenylating and destabilizing elements. UAUUUAUU elements provided a strong deadenylating activity that was dependent on the activation of the zygotic genome, with a weaker effect on mRNA decay. This suggest that the dynamics of mRNA decay may depend on the pathway promoting deadenylation. On the same mRNA, RESA identified multiple sequence elements with antagonistic effects. For example, the *zorba* mRNA contains adjacent stabilizing poly-U motifs and destabilizing miR-430 target sites. Combining these elements on the same mRNAs may provide differential temporal regulation. We propose that these transcripts are first stabilized by maternally provided poly-U binding proteins and later deadenylated and degraded by the zygotic mode of decay triggered by the activation of miR-430 or UAUUUAUU elements (Audic et al, 1997; Voeltz et al, 1998; Giraldez et al, 2006; Wu et al, 2006; Bazzini et al, 2012). Initial stabilization is likely coupled with active cytoplasmic polyadenylation, which would increase mRNA translation efficiency early in embryonic development (Subtelny et al, 2014). We find a strong correspondence between the motifs identified by RESA and those recently reported in zebrafish by Rabani et al, 2017. We further characterized the sequence contexts that influence the strength of the regulation associated with these elements. We identified an antagonistic activity between poly-U and CCUC sequences. Indeed, transcriptome-wide analysis of 3’-UTR mediated stability reveal a global tendency of 3’-UTR sequences (rich in U) to promote stabilization of the mRNA, with destabilizing islands that contain three main elements: miRNAs, CCUC domains, and UAUUUAUU motifs. Furthermore, the fact that deadenylation and decay are initially uncoupled during embryogenesis allows us to define elements that preferentially affect mRNA stability (poly-U and CCUC) (Stoeckius et al, 2014), elements that mainly affect mRNA deadenylation (UAUUUAUU-ARE) and elements that induce both, such as miR-430. ELAV-like proteins (HuR) have been shown to stall ARE-mediated decay downstream of deadenylation (Fan et al, 1998; Peng et al, 1998) and miRNAs (Kundu et al, 2012). The favorable sequence contexts for HuR and polyC-BP are similar to their binding motif, and show a broader accumulation of iCLIP signal across the binding sites, suggesting that favorable context sites are bound by multiple proteins.

Using iCLIP we characterized the binding motifs of multiple RBPs identified in the interactome capture. We observed that similar motifs are recognized by multiple RBPs, which are associated with similar regulatory activities. While a high level of functional redundancy could be part of a robust developmental system, these RBPs might also recruit other proteins to achieve a specific regulatory response. Functional redundancy impairs our ability to genetically dissect their activity, an issue further complicated by the existence of multiple homologs for each RBP. Nevertheless, the genetic elimination of the regulatory elements identified in RESA clearly reveals the importance of these sites in the regulation of endogenous mRNA levels (Figure 6H, I). These regulatory elements are enriched at the primary sequence level, raising a conundrum about the origin of the enrichment of poly-U and depletion of CCUC sequences in the 3’-UTRs. During evolution, 3’-UTRs might have evolved to recruit RBPs that promote mRNA stability. Alternatively, RBPs were adapted to bind non-coding sequences. While our study leaves this question open, the high density of binding we observed for U-rich sequences within 3’-UTRs emphasizes their central role in regulating RNAs.

We integrated these regulatory activities into a model using machine learning. Our model was able to predict regulatory activity across the RESA library of reporters for maternal and zygotic signals that mediate mRNA stability. We also identify sequence motifs associated with lower polyadenylation status in the mRNA (namely miR-430 and UAUUUAUU) dependent on the zygotic program. The stability of endogenous mRNAs was less accurately predicted for endogenous mRNAs (r=0.29) than for the RESA 3’-UTR reporter library (per transcript Pearson r=0.82, Supp. Figure 9A). Our model is solely based on the primary sequence of the 3’-UTR, and its lower predictive power in endogenous transcripts is consistent with recent studies (Rabani et al, 2017). While using the same reporter backbone for all fragments tested in RESA allowed us to eliminate any bias due to different coding sequences, other factors not probed by RESA are regulating mRNA fate. Notably, the codon bias of the coding-sequence of mRNAs influences their stability (Hanson et al, 2017), specifically during the MZT (Bazzini et al, 2016; Mishima et al, 2016). Also, our RESA libraries are synthesized *in vitro* and therefore lack any RNA modification. Modifications such as m6A have been proposed to help the clearance of maternal mRNAs (Zhao et al, 2017), yet it is unclear whether m6A containing mRNAs are specifically stabilized upon loss of m6A reader proteins (Kontur et al, 2017). Finally, RNA structure analysis has identified long range RNA-interactions that are currently not being integrated in the RESA reporters due to the limited size of the fragments tested (Lu et al, 2016; Aw et al, 2016; Sharma et al, 2016). Explaining the decay dynamics of all mRNAs will require the integration of the regulatory activities found in the 3’-UTRs with additional elements in the mRNA, namely codons, RNA structure and modifications to achieve a global prediction of mRNA dynamics.

## Author contributions

CMT, VY, CEV, DC, MAM and AJG conceived the project. CMT, VY, DC, SL, RC, MS and HDC performed the experiments. CEV, MAM, PO and RC performed computational analysis of the data. CMT, CEV, VY and MTL performed RESA analysis. DC, SL and RC performed interactome capture. MAM performed random forest analysis, model-based motifs identification and reporter mRNA predictions. DC, SL, and CEV performed iCLIP and data analysis. CMT, VY and MS performed reporter analysis. PO performed FIRE analysis. MAM, JDB, HDC and CEV performed *trip10* reporter analysis. CMT, VY, DC, CEV, MAM, PO, MS, RC and AJG interpreted the results. AJG supervised the project, with the contribution of ST and TW. CEV, CMT and VY drafted the manuscript. CEV, MAM and AJG wrote the manuscript with input from the other authors.

### Acknowledgements

We thank Karen Bishop for technical help, and Ariel Bazzini, Miguel Moreno-Mateos and all the members of the Giraldez laboratory for intellectual and technical support. Dr. Matthias Hentze and Dr. Alfredo Castello for providing the original interactome capture protocol. Dr. Valerie Tornini for manuscript editing. The Swiss National Science Foundation (grant P2GEP3_148600 to CEV), NIH grants NHGRI (2R01HG003219) and NHGRI (1R01HG009065) (PO and ST), the Deutsche Forschungsgemeinschaft (DFG) (MS fellowship), the Eunice Kennedy Shriver National Institute of Child Health and Human Development-NIH grant K99HD071968 (DC), HHMI Faculty Scholars program (AJG), NIH grants R21 HD073768, R01 HD073768, R01 GM102251, and R35 GM122580 (AJG) supported this work. The Yale Scholars Program and Whitman fellowship funds provided by E. E. Just, Lucy B. Lemann, Evelyn and Melvin Spiegel, The H. Keffer Hartline and Edward F. MacNichol, Jr. of the Marine Biological Laboratory in Woods Hole, MA to AJG. The research of AJG was supported in part by a Faculty Scholar grant from the Howard Hughes Medical Institute and the Simons Foundation.

## Methods

### Timecourse sample collection and RNA-seq library preparation

For the large developmental gene expression time course, synchronously developing embryos were collected every 30 minutes (between 0 hours per fertilization, hpf, and 5 hpf) and at 6 hpf and 8 hpf. Ten embryos were collected for each time point in duplicates. To allow for appropriate normalization of fold changes, yeast total RNA (roughly 5% of estimated zebrafish total RNA) was spiked into Trizol (Invitrogen, USA) prior to RNA extraction according to manufacturer’s instructions (Invitrogen, USA). After extraction, RNA was subjected to poly(A)-selected RNA-seq library preparation (TruSeq mRNA, Illumina, USA) and ribosomal RNA-depleted total RNA-seq library preparation (TruSeq Total RNA, Illumina, USA) according to manufacturer’s instructions. For Pol II inhibition, *α*-amanitin was obtained from Sigma Aldrich and resuspended in nuclease-free water. Dechorionated embryos were injected with 0.2ng of *α*-amanitin at one-cell stage.

### RNA-Seq analysis: Developmental modes and their contribution to decay

Raw reads were mapped to the zebrafish Zv9 and yeast R64-1-1 genomes using STAR (Dobin et al, 2013) version 2.4.2a with the following non-default parameters: *╌alignEndsType ╌Local ╌outFilterMultimapNmax 100 ╌seedSearchStartLmax 30 ╌sjdbScore 10*. Genomic sequence indices for STAR were built including exon-junction coordinates from Ensembl r78 (Aken et al, 2017). Read counts per gene were computed by summing the total number of reads overlapping at least by 10 nucleotides the gene annotation, including only uniquely mapped reads in the genome. Per gene annotation was obtained by concatenating all Ensembl isoforms together. To determine significantly over- and under-expressed genes, gene read counts were compared using DESeq2 (Love et al, 2014). Genes below 1 count in all replicates in either condition were excluded from the analysis. Total gene counts for all yeast genes were summed and the ratios between conditions were input as Factors to DESeq2. To get DE genes, the *results* function was used with *pAdjustMethod="fdr"* and *independentFiltering=FALSE* options. To categorize decay genes in *Maternal*, *Zygotic: miR-430 dependent* and *Zygotic: miR-430 independent* modes, conditions 2 hpf, 2 hour (h) *α*-amanitin, 6 hpf, 6 h *α*-amanitin and 6 h LNA miR-430 were used. First, decaying genes were selected as significantly higher at 2 hpf vs. 6 hpf and not significantly higher at 6hpf vs. 6h *α*-amanitin. Second, among decaying genes, *Zygotic: miR-430 dependent* were selected as significantly higher at 6h LNA miR-430 vs. 6 hpf. Remaining genes were then split into two modes. If only 6 h *α*-amanitin was significantly higher than 6 hpf, genes were classified as *Zygotic: miR-430 independent*. If only 2 h *α*-amanitin was significantly higher than 6 h *α*-amanitin, genes were classified as *Maternal*. If a gene could be classified in both lists, the most significant program was prioritized, i.e. the lowest P-value determined the program. To calculate the contribution of each program, *Maternal*, *Zygotic: miR-430 dependent* and *Zygotic: miR-430 independent*, log2 fold-changes as determined by DESeq2 for *Maternal program*, (2 h *α*-amanitin vs. 6 h *α*-amanitin), *Zygotic: miR-430 dependent* (6 hpf vs. 6 h LNA miR-430) and *Zygotic: miR-430 independent* (6 hpf vs. 6 h *α*-amanitin) were used. Because this analysis used the 3 fold-changes corresponding to each program and was not limited to the single statistical test used to determine the main program, fold-changes with biologically non-expected sign were set to not contribute: the three ratios above for *Maternal*, *Zygotic: miR-430 dependent* and *Zygotic: miR-430 independent* program fold-changes that were positive, negative and negative, respectively, were set to zero. The *α*-amanitin treatment also blocking *miR-430* expression, the *Maternal* and *Zygotic: miR-430 independent* modes were first normalized together to 1 and then within *Zygotic* modes, both *miR-430 dependent* and *independent* fold-changes were normalized.

### In situ hybridization

*In situ* hybridization was performed as in Thisse et al, 2008 with 20ng of DIG-labeled RNA probe per 200μL hybridization reaction. RNA probes were synthesized from T7 promoter polymerase overhang on the reverse oligo. Oligo sequences used to amplify transcript regions are listed below: In situ probes: org_For: TGACTGACCAGTGCAACTACG org_Rev: AAACAGCAAATCGAGAAGCAA trip10_For: ATGGACTGGGGAACTGAGCTTTG trip10_Rev: GAGAACCATAGAGTCATTCCTCG dnajc5ga_For: ATCGCTGTACGCTTTCAAGG dnajc5ga_Rev: AAAACCCACTTCCCTTCTGG

### Reporter validation by qRT-PCR

Validation sequences were cloned into PCS2+ vector downstream of GFP using XbaI and XhoI restriction sites. RNA was in vitro synthesized using Sp6 Message Machine (ThermoFisher AM1340) from NotI linearized plasmids. Zebrafish embryos were injected with 4pg of each reporter and dsRED control mRNAs and collected at 2hpf and 8hpf. Total RNA (250ng) was reverse transcribed using SuperScript III Kit (Invitrogen #18080-051) with random hexamers. cDNA was diluted 1:20 and used in 10uL reaction (5μL SYBR green master mix (ThermoFisher #4472908), 0.5μL 10μM Forward and Reverse primer mix, 1μL 1:20 diluted cDNA, 3.5μL water). A common forward primer complementary to GFP - CATG- GTCCTGCTGGAGTTCGTGAC and a reporter-specific reverse primer (UACA - GTACATGAGACTCAATCACTGCTC, tdrd7 -TTCGAAACACCATGGATGCTTCTC). DsRED was amplified with Forward GAAGGGCGAGATCCACAAG and Reverse GGACTTGAACTCCACCAGGTA primers. Biological and technical triplicates were performed for each sample and relative expression with ΔΔCT method was measured using ViiA 7 Software v1.2.2 with dsRED mRNA as a reference control.

### RESA reporter library construction

The transcriptome-based reporter library was generated with a previously constructed Illumina RNA sequencing library Bazzini et al, 2012 by overlap-extension PCR with primers mapping to the SP6 promoter and downstream of the SV40 polyadenylation site (Yartseva et al, 2017). Five separate extension reactions were performed (5 cycles of 98C, 61C, and 72C), then pooled and used as a template for further amplification (48 separate PCR reactions of 10 cycles each). PCR products were pooled and concentrated by MinElute PCR Purification Kit (Qiagen), followed by PAGE purification. Following *in vitro* transcription, reporter mRNA was injected into ~75 embryos at 100ng/μL (1 nl). Total RNA was extracted from embryos using TRIzol reagent according to manufacturer’s instructions. Reverse transcription was performed with a reporter-specific primer (CATCAATGTATCTTATCATGTCTGGATC; SuperScript III (Invitrogen)). The final Illumina library was prepared using the following primers: 5’-AATGATACGGCGACCACCGACAGGTTCAGAGTTCTACAGTC-CGACGATC-3’ and 5’-CAAGCAGAAGACGGCATACGAGAT(barcode)GTGACTGGAGTTCAGACGTGTGCTCTTCCGATCT-3’. PCR reactions were pooled (5 reactions (17 cycles each; Phusion (NEB))) and purified by PAGE. The targeted library was prepared as described previously (Yartseva et al, 2017).

T1 5’ (Sp6 promoter, GFP coding sequence, and 5’ Illumina adaptor) GCTTGATTTAGGTGACACTATAGAATACAAGCTACTTGTTCTTTTTGCAGGATCCATGGTGAGCAAGGGCGAGGAGCTGTTCACCGGGGTGGTGCCC ATCCTGGTCGAGCTGGACGGCGACGTAAACGGCCACAAGTTCAGCGTGTCCGGCGAGGGCGAGGGCGATGCCACCTACGGCAAGCTGACCCTGAAGT TCATCTGCACCACCGGCAAGCTGCCCGTGCCCTGGCCCACCCTCGTGACCACCCTGACCTACGGCGTGCAGTGCTTCAGCCGCTACCCCGACCACAT GAAGCAGCACGACTTCTTCAAGTCCGCCATGCCCGAAGGCTACGTCCAGGAGCGCACCATCTTCTTCAAGGACGACGGCAACTACAAGACCCGCGCC GAGGTGAAGTTCGAGGGCGACACCCTGGTGAACCGCATCGAGCTGAAGGGCATCGACTTCAAGGAGGACGGCAACATCCTGGGGCACAAGCTGGAGT ACAACTACAACAGCCACAACGTCTATATCATGGCCGACAAGCAGAAGAACGGCATCAAGGTGAACTTCAAGATCCGCCACAACATCGAGGACGGCAG CGTGCAGCTCGCCGACCACTACCAGCAGAACACCCCCATCGGCGACGGCCCCGTGCTGCTGCCCGACAACCACTACCTGAGCACCCAGTCCGCCCTG AGCAAAGACCCCAACGAGAAGCGCGATCACATGGTCCTGCTGGAGTTCGTGACCGCCGCCGGGATCACTCTCGGCATGGACGAGCTGTACAAGTCCG GACTCAGATCTAAGCTGAACCCTCCTGATGAGAGTGGCCCCGGCTGCATGAGCTGCAAGTGTGTGCTCTCCTGACTCGAGGATCTACACTCTTTCCC TACACGACGCTCTTCCGATCT

T2 3’ (Illumina adaptor and SV40 polyadenylation signal) AGATCGGAAGAGCACACGTCTGAACTCCAGTCACTCTAGAACTATAGTGAGTCGTATTACGTAGATCCAGACATGATAAGATACATTGATGAGTTTGGACAAACCACAACTAGAATGCAGTGAAAAAAATGCTTTATTTGTGAAATTTGTGATGCTATTGCTTTATTTGTAACCATTATAAGCTGCAATAAACAAGTTAACAACAACAATTGCATTCATTTTATGTTTCAGGTTCAGGGGGAGGTGTGGGAGGTTTTTTAATTC

### Genetic deletion of the trip10 3’-UTR region

CRISPR-mediated mutagenesis was performed as described in Moreno-Mateos et al, 2015. Standard PCR with Taq enzyme was used for genotyping with For-CATGAAAGCCACGTCGATAA and Rev-CAAATGCAAAACAAACACTCG and annealing temperature of 61°C.

### RESA profiles

The Illumina TruSeq index adaptor sequence was first trimmed from raw reads (i) for transcriptomic library by aligning its sequence, requiring 100% match of the first five base pairs and a minimum global alignment score of 60 (Matches: 5, Mismatches: −4, Gap opening: −7, Gap extension: −7, Cost-free ends gaps) or (ii) for targeted library using Skewer (Jiang et al, 2014). Reads were then mapped to the zebrafish Zv9 genomes using STAR (Dobin et al, 2013) version 2.4.2a with the following non-default parameters: *╌alignEndsType EndToEnd ╌outFilterMultimapNmax 100 ╌seedSearchStartLmax 30 ╌sjdbScore 2*. Genomic sequence indices for STAR were built including exon-junction coordinates from Ensembl r78 (Aken et al, 2017). Individual RESA profiles for each transcript (transcriptomic library) or UTR (targeted library) are positional read coverage. Coverage was calculated using pooled biological replicates by incrementing each position in the profile from first to the last overlapping positions of reads that were sense to the transcript/UTR. For the targeted library, sequenced with paired-end reads, both reads were merged into a single fragment, and every position of this fragment was used to increment the coverage. Profiles were then normalized to CPM (Count Per Million) using total counts of all transcript/UTR profiles per sample.

### FIRE analysis

#### Regulatory categories of transcriptome segments

To evaluate the regulatory effect of short RNA elements, RESA profiles were split into fixed-length sequence segments (30 and 100 nucleotides long for the transcriptomic and targeted libraries respectively) with 33% overlaps (Oikonomou et al, 2014). For every segment and each library, a score was calculated by averaging over the length of the segment the log-transformed frequency of read counts observed at each base. When comparing between two libraries at different time points (e.g. 2hpf and 6hpf) or treatments (e.g. wild type and LNA-treated), the difference of the scores of a given segment in the two libraries is transformed into a Z-score from the distribution of the all differences (e.g. *z(WT_2hpf; WT_6hpf)* is the Z-score of the difference between wild type at 2hpf and 6hpf). Z-scores can be used to evenly compare differences between different pairs of libraries. To determine whether a given sequence segment is modulated by either a maternal, zygotic, *miR-430* or stabilizing regulatory effect, the following strategy was used. For each segment, changes in sequencing coverage between the following five pairs of libraries were considered: (1) wild type at 2 hpf and 6 hpf (2) *α*-amanitin treated at 6 hpf and wild type at 6 hpf (3) LNA-430 treated at 6 hpf and wild type at 6 hpf (4) *α*-amanitin treated at 6 hpf and wild type at 2 hpf (5) LNA-430 treated at 6 hpf and wild type at 2 hpf. For each pair, Z-scores are computed for every segment as described above. Functional categories were then assigned to each sequence segment based on the values of these Z-scores. For the destabilizing regulatory categories, only segments with a sequencing score at 2 hpf above threshold (segment score > 2) were considered to ensure that the sequence segment was present at the early stage. The maternal category were defined as: *z(WT_2hpf; WT_6hpf)<-z^th^ and z(WT_2hpf; Am_6hpf)<-z^th^ and z(WT_2hpf; LNA_6hpf)<-z^th^*, for a permissive threshold of *z^th^=0.5*. This pattern corresponds to a decaying number of reads at 6 hpf with respect to 2 hpf for the wild type that is not alleviated by *α*-amanatin or LNA-430 treatments. The zygotic category was defined as: *z(WT_2hpf; WT_6hpf)<-z^th^ and z(WT_6hpf; Am_6hpf)>z^th^ and z(WT_2hpf; LNA_6hpf)<-z^th^*, which corresponds to a decaying number of reads at 6 hpf for the wild type that is alleviated by *α*-amanatin but not by LNA-430 treatment. The miR-430 category was defined as: *z(WT_2hpf; WT_6hpf)<-z^th^ and z(WT_6hpf; Am_6hpf)>z^th^ and z(WT_6hpf; LNA_6hpf)>z^th^*, which corresponds to a decaying number of reads at 6 hpf for the wild type that is alleviated by both *α*-amanatin and LNA-430 treatments. Corresponding categories were defined for stabilizing modes of regulation by inverting the inequality signs in the above boolean statements, for example maternally stable segments are defined as: *z(WT_2hpf; WT_6hpf)>z^th^ and z(WT_2hpf; Am_6hpf)>z^th^ and z(WT_2hpf; LNA_6hpf)<z^th^*. Given this scheme, a particular sequence segment can be assigned to multiple categories. To resolve this discrepancy *P*-value associated with each category were calculated. The three Z-scores used to define a category correspond to three *P*-values combined using Fisher’s method. Each segment was assigned to the category with lowest p-value. This process was repeated for both sets of replicates of the libraries and final categories were assigned to the sequence segment if the categories for the two replicates were both decaying or are both stabilizing. Benjamini-Hochberg corrected q-values were then calculated for each sequence and the number of segments assigned to various categories shown in Supp. Figure 3A (q-value<0.05).

### Post-transcriptional Regulatory Element Discovery

The identification distinct stabilizing and destabilizing categories of sequences enables *de novo* discovery of short motifs that are significantly informative of the different modes of regulation. Here, an updated version of FIRE, a computational framework for the discovery of regulatory elements (Elemento et al, 2007) was used to systematically explore the space of linear RNA elements within tens of thousands of sequences that fall into multiple categories simultaneously. N-fold cross-validation option was introduced in FIRE, whereby the dataset is partitioned into N sets, N-1 parts are used as the training set for motif discovery and 1 set is left aside as a test set to evaluate the results. From the motifs discovered on the training set, only ones that exhibit significant mutual information in the test set (Z-score>2) and have a pattern of over- and under-representation across categories that is similar to the representation pattern in the training set (correlation>0.5) were kept. The motif discovery was repeated N times with each of the N sets used once as the test set. The resulting motifs were combined and similar motifs were filtered out by means of sequence similarity and conditional information (Elemento et al, 2007; Oikonomou et al, 2014). The introduction of cross-validation avoids data over-fitting and produces a shorter, but more robust list of regulatory elements.

Groups of RNA trans-acting factors had binding sites that are highly similar with respect to nucleotide composition and are significantly shorter than the binding sites of transcription factors. In combination with the greedy nature of motif discovery algorithms this often results in degenerate and often uninterruptible motif representations when analyzing large libraries for linear RNA regulatory elements. Here, average degeneracy allowed for the regular expression of each motif was capped, so that it is optimized for maximal mutual information while sustaining a well-defined sequence profile.

FIRE with zebrafish specific files, so that motif discovery can be applied to identify DNA and RNA regulatory elements for zebrafish expression data and updated release of FIRE and the related files can be downloaded at: https://tavazoielab.c2b2.columbia.edu/XXXX/

The categorized sequence segments from the RESA libraries are further analyzed for motif discovery. A more permissive list of sequences (*P*-value<0.05; 20,111 segments for the transcriptomic library, 3,090 and 2,642 for the targeted total RNA and poly(A)-selected library respectively) were split into the categories described above (stabilizing and destabilizing maternal, zygotic and miR-430). Within each category, segments are sorted based on their *P*-value and further grouped into equally sized bins (900 and 400 segments per bin for transcriptomic and targeted libraries respectively). *De novo* motif discovery with 3-fold cross-validation on these categories was performed. FIRE revealed a total of 45 RNA motifs for the transcriptomic library and 38 motifs for the targeted library that were significantly informative of the observed regulatory categories. The most significant of these results are shown in Figure 4.

Over representation (yellow) and under representation (blue) patterns are shown for each discovered motif within each category of sequence segments. Mutual information values, Z-scores associated with a randomization-based statistical test and robustness scores from a three-fold jackknifing test are also shown (Elemento et al, 2007).

### Reporter validation zc3h18 (poly-C motif)

Oligos:

wt-F: tcgagCTGCCTCCTCCACCTCCTCCTCTAGATCCTCCAGt

wt-R: ctagaCTGGAGGATCTAGAGGAGGAGGTGGAGGAGGCAGc

mut-F: tcgagCTGCGTCGTCCACGTCGTCCTCTAGATCCTCCAGt

mut-R: ctagaCTGGAGGATCTAGAGGACGACGTGGACGACGCAGc

Forward and reverse oligos were annealed and cloned into pCS2-GFP (XhoI-XbaI), followed by linearization with NotI and *in vitro* transcription from Sp6 promoter (mMessage mMachine (Ambion)). mRNA was co-injected with mRNA encoding DsRed as described previously.

### Hire-PAT reporter assay

Fam116b (ARE motif)

Forward oligo: 5’-CTCGAGTTTTCCCATGATGAAATTAGG-3’

Reverse oligo: 5’-TCTAGACCGATGCCATAGTGTTCTTG-3’

A 107 nucleotide fragment of the 3’-UTR of *fam116* was cloned into pCS2-GFP (XhoI-XbaI), followed by linearization with NotI and *in vitro* transcription from Sp6 promoter (mMessage mMachine (Ambion)). mRNA was co-injected with mRNA encoding DsRed as described previously. Total RNA was extracted from 6 hpf embryos using TRIzol reagent according to manufacturer instructions. One microgram of total RNA was used for the poly(A) tail length assay according to manufacturer instructions (Affymetrix Poly(A) Tail-Length Assay Kit). Poly(G/I) tailed cDNA was amplified using the a *fam116*-specific forward primer (5’-CTGAAGTGCACAGAAGAACCAC-3’) and a FAM-labeled universal reverse primer (FAMGGTAATACGACTCACTATAGCGAGACCCCCCCCCCTT). Fragment analysis was performed in a 3730XL 96-Capillary Genetic Analyzer and output was analyzed using GeneMarker software (Softgenetics).

### Interactome Capture

Wild type zebrafish embryos were irradiated at 4 h pf with UV at 254 nm for 4 minutes in a Spectrolinker XL-1000 UV crosslinker. During this time, the embryos were covered with water in 6-well multiwell plate on ice and at a distance from the UV source of ~9 cm. Immediately after cross-linking, the embryos were collected in 1.5 ml tubes and flash frozen in liquid nitrogen. *α*-amanitin-injected embryos were processed in the same way. The –UV control group of embryos were not UV-irradiated prior to collection. In total, 3 replicates of +UV wild-type embryos, +UV *α*-amanitin-injected embryos, and 3 replicates of –UV embryos were processed. Each replicate contains around 4,000 embryos. Interactome capture was conducted as described in Castello et al, 2012. Embryos were lysed in Lysis/Binding Buffer (20 mM Tris-HCl (pH 7.5), 500 mM LiCl, 0.5% LiDS, 1 mM EDTA, 5 mM DTT) and the poly(A)^+^ mRNAs with the associated cross-linked proteins were pulled-down with oligo d(T)^25^ Magnetic Beads (NEB). Next, the pellets of oligo d(T)^25^ beads were sequentially washed with buffers with decreasing concentration of LiDS and LiCl (Wash Buffer I (20 mM Tris-HCl (pH 7.5), 500 mM LiCl, 0.1% LiDS, 1 mM EDTA, 5 mM DTT); Wash Buffer II (20 mM Tris-HCl (pH 7.5), 500 mM LiCl, 1 mM EDTA, 5 mM DTT); Low Salt Buffer (20 mM Tris-HCl (pH 7.5), 200 mM LiCl, 1 mM EDTA; Elution Buffer (20 mM Tris-HCl (pH 7.5), 1 mM EDTA)). After washes, the pellets of beads were frozen, awaiting further processing. Crosslinked material was rinsed three times with wash buffer (150 mM NaCl, 50 mM Tris, pH 7.4) and then denaturated for 30 minutes with urea (8 M) in 0.1 M Tris, pH 7.4, 1 mM DTT before alkylation and on-bead pre-digestion with Endoproteinase LysC (Wako Chemicals USA, Inc.). After incubation for 3 hours, samples were diluted 4-fold with ammonium bicarbonate (25 mM) and further digested with trypsin (Promega) overnight. Digestions were stopped by addition of trifluoroacetic acid (TFA, 1 µL), and the resulting peptides were loaded and desalted on C18 Stage Tips.

### LC-MS/MS analysis

Peptides were eluted from C18 Stage Tips with 60 μL of elution buffer (80% acetonitrile and 0.1% formic acid), and samples were dried down to 5 μL in a vacuum centrifuge. Peptides were then subjected to reversed phase chromatography on an Easy nLC 1000 system (Thermo Fisher Scientific) using a 50-cm column (New Objective) with an inner diameter of 75 μm, packed in-house with 1.9 μm C18 resin (Dr. Maisch GmbH). Peptides were eluted with an acetonitrile gradient (5–30% for 95 min at a constant flow rate of 250 nL/min) and directly electrosprayed into a mass spectrometer (Q Exactive; Thermo Fisher Scientific). Mass spectra were acquired on the spectrometer in a data-dependent mode to automatically switch between full scan MS and up to 10 data-dependent MS/MS scans. The maximum injection time for full scans was 20 ms, with a target value of 3,000,000 at a resolution of 70,000 at m/z = 200. The ten most intense multiple charged ions (z ≥ 2) from the survey scan were selected with an isolation width of 3Th and fragmented with higher energy collision dissociation (HCD) with normalized collision energies of 25. Target values for MS/MS were set to 100,000 with a maximum injection time of 160 ms at a resolution of 17,500 at m/z = 200. To avoid repetitive sequencing, the dynamic exclusion of sequenced peptides was set to 35 s.

### MS data analysis

MS and MS/MS spectra were analyzed using MaxQuant (version 1.3.8.1), utilizing its integrated ANDROMEDA search algorithms. Scoring of peptides for identification was carried out with an initial allowed mass deviation of the precursor ion of up to 6 ppm for the search for peptides with a minimum length of six amino acids. The allowed fragment mass deviation was 20 ppm. The false discovery rate (FDR) was set to 0.01 for proteins and peptides. Peak lists were searched against a local database for human proteome. Maximum missed cleavages were set to 2. The search included carbamidomethylation of cysteines as a fixed modification and methionine oxidation and N-terminal acetylation as variable modifications. The final list of proteins was curated to remove duplicates and retain only proteins with at least two peptides and at least one of them unique.

### Whole-embryo lysate proteomics

4 hpf embryos were collected in triplicates in deyolking buffer (55 mM NaCl, 1.8 mM KCl, 1.25 mM NaHCO^3^) and pipetted up and down to disrupt the embryos. Cells were pelleted by centrifuging 30 seconds at 300 x g. The pellet of cells was then washed twice with wash buffer (110 mM NaCl, 3.5 mM KCl, 2.7 mM CaCl^2^, 10 mM Tris-HCl pH8.5), followed by flash freezing in liquid nitrogen. Samples were further processed and analyzed by mass spectrometry as described earlier for the Interactome capture.

## Data Availability

All raw mass spectrometry data files from this study are available at ProteomeXchange (XXXX) repositories.

### Gene Ontology enrichment

Enrichment in the molecular function category was tested by comparing the proteins found in the interactome against all the proteins identified in the whole-embryo lysate of wild-type and *α*-manitin-injected embryos at 4 hpf. Gene Ontology annotation for these proteins was downloaded from ENSEMBL using BioMart. For each GO term, the number of proteins identified in the interactome with this GO term was normalized by the total number of proteins in all GO terms. The same calculation was conducted with the proteins in the input. The odds ratio for a given GO term is the result of dividing the value obtained from interactome proteins by value obtain from the input. Only GO terms present in both samples and with a log2 odds ratio >2.5 or < -2.5 were retained for further analysis.

### RBP Cloning

To clone each selected RBP into pCS2+, specific primers were designed using Primer3 (Rozen & Skaletsky, 1998) that also incorporate the corresponding restriction enzyme. The 5’ primers also contain the sequence encoding the FLAG tag in frame with the downstream RBP. Next, the cDNA of each RBP was retrotranscribed from total RNA isolated from zebrafish embryos at mixed stages using SuperScript III Reverse Transcriptase (Invitrogen) and amplified with Phusion^®^ High-Fidelity DNA Polymerase (NEB). Next, each PCR amplicon was cut with the appropriate set of restriction enzymes and ligated to a linearized pCS2+ vector with T4 DNA ligase (NEB). Finally, the resulting clones were linearized and *in vitro* transcribed to capped mRNA using the mMESSAGE mMACHINE SP6 Transcription Kit (Ambion). Oligos are listed in Supp. Table 2.

### iCLIP library cloning

The iCLIP protocol described in Huppertz et al, 2014 has been modified and adapted to zebrafish.

### Sample collection

For each given RBP, zebrafish embryos at 1-cell were injected with 1 nl of the corresponding capped mRNA at 0.1 μg/μL. Sphere stage embryos (~4 hpf) were cross-linked with UV at 254 nm for 4 minutes. Plates of embryos were deposited over a tray of ice during cross-linking and immediately collected into 1.5 ml eppendorf tubes, flash frozen in liquid nitrogen, and stored at -80°C until processing. Around 200 embryos were collected per replicate.

### Lysis and pull-down

Embryo samples were lysed with 1 ml of lysis buffer (50 mM Tris–HCl, pH 7.4, 100 mM NaCl, 1% Igepal CA-630, 0.5% sodium deoxycholate, 0.1% SDS, 1/100 volume of Protease Inhibitor Cocktail Set III (Calbiochem), 1/1000 volume of SUPERase•In RNase Inhibitor (Ambion)), and vortexed. 4 μL of Turbo DNAse (Ambion) were added to each tube, followed by 10 μL of RNAse I (Invitrogen) diluted to 1:200 in PBS. Samples were incubated in the thermomixer for 3 minutes at 37ºC and 1,100 rpm. Immediately after, 5 μL of RNase Inhibitor were added, and the samples were incubated on ice for at least 3 minutes. Nest, samples were centrifuged at 4ºC for 10 minutes at 14,000 rpm. In the meantime, Anti-FLAG^®^ M2 Magnetic Beads (Sigma) were prepared at a ratio of 20 μL of beads per 200 embryos. Beads were washed 3 times with 800 μL of lysis buffer, avoiding bubbles. Next, the supernatant was transferred to the pellet of antibody with beads and incubated for 2 hours at 4ºC in an orbital rocker. For Ddx6 iCLIP, Ddx6 was pulled-down using a specific antibody (Abcam, ab40684) and Dynabeads Protein G (Invitrogen) in three replicates of ~1,000 embryos each. After, pull-down, beads were washed twice with High Salt buffer (50 mM Tris–HCl, pH 7.4, 1 M NaCl, 1 mM EDTA, 1% Igepal CA-630, 0.1% SDS, 0.5% sodium deoxycholate) and 3 times with PNK buffer (20 mM Tris–HCl, pH 7.4, 10 mM MgCl^2^, 0.2% Tween-20), 800 μL each time.

### 5’ labeling of RNA fragments

4 μL of the labeling reaction (0.2 μL PNK, 0.4 μL gamma-^32^P-ATP, 0.4 μL 10x PNK buffer, 2.8 μL H2O, 0.2 μL RNase inhibitor) were added to the pellet of washed beads prior incubation of 5 minutes at 37ºC and 1,100 rpm in the thermomixer. The pellets were washed one time with 500 μL PNK buffer.

### NUPAGE electrophoresis and membrane transfer

30 μL of NUPAGE loading sample buffer with reducing agent were added to the beads and incubated 10 minutes at 80 ºC, shaking at 1,110 rpm. Next, the supernatant was loaded into a precast NUPAGE 4-12% gradient gel in the Bolt system (Invitrogen). The electophoresis ran for 50 minutes at 180 V. After the run, the riboprotein complexes from the gel were transferred to a nitrocellulose membrane using the iBlot system (Invitrogen). After the transfer, the membrane was exposed to a storage phosphorimager plate for 3 hours at -20 ºC.

### Proteinase K digestion

The membrane sections containing the ribo-protein complexes according to the scan of the storage phosphorimager screen were cut and added to a 2 ml tube and with 200 μL of Proteinase K mix (Tris-HCl 100 mM, NaCl 50 mM, EDTA 10 mM, Proteinase K). The samples were incubated 20 minutes at 37 ºC and 1,100 rpm and an additional 200 μL of Proteinase K buffer with 7 M urea were added, followed with an additional incubation of 20 minutes at 37 ºC and 1100 rpm. The RNA fragments were extracted from the membrane with 400 μL of phenol-chloroform and vigorous shaking for 10 minutes at room temperature. The liquid was transferred to a new tube and centrifuged 10 minutes at maximum 14,000 rpm. The aqueous phase was collected and the RNA was precipitated with 40 μL of sodium acetate 3 M, 0.7 μL glycogen and 1 ml of 96% ethanol.

### 3’-end repair (dephosporylation)

The repair reaction mix (2 μL of 10x PNK pH 6.5 buffer, 0.5 μL polynucleotide kinase (NEB), 0.5 μL RNase Inhibitor, 15 μL water) was added to the purified pellet of RNA and incubated for 20 minutes at 37ºC, followed by RNA precipitation.

### 3’ pre-adenylated adaptor ligation

The RNA was resuspended with of 10 μL water plus the ligation reaction mix (20 pmoles pre-adenylated DNA adapter, 2 μL 10x T4 RNA Ligase 2 truncated K227Q reaction buffer (NEB), 1 μL 200 U T4 RNA Ligase 2 truncated K227Q (NEB), 1 μL of RNaseOUT (Invitrogen), 3 μL of 50% PEG8000 and 4 μL of water). Ligation reactions were incubated at 16 °C overnight. Each sample within a replicate of the same RBP was ligated with an adaptor with different barcodes to allow sample pooling in the subsequent steps. The sequence of the 3’ adaptor used was: rApp-NNcacaACTGTAGGCACCATCAAT-ddC, where lowercase represents one of the custom barcodes.

### Size fractionation

To remove the unligated adaptor, all replicate samples were combined and loaded in a 15% denaturing polyacrilamide-urea gel. After running, the gel was exposed in a storage phosporimager screen and kept at -20 °C. Next, gel slices containing the ligated RNA were cut and RNA was extracted and precipitated.

### Retrotranscription

The purified RNA was resuspended in 8 μL of water with 1 μL RT iCLIP primer (0.5 pmol/μL of the oligo -(Phos) NNNNAGATCGGAAGAGCGTCGTGTAGGGAAAGAGTGTAGATCTCGGTGGTCGC-(SpC18)-CACTCA-(SpC18)-TTCAGACGTGTGCTCTTCCGATCTATTGATGGTGCCTACAG), 1 μL dNTP mix (10 mM) and incubated for 5 minutes at 70 ºC. Next the retrotrasncription reaction mix was added: 4 μL H2O, 4 μL 5x First Strand Buffer, 1 μL 0.1 M DTT, 0.5 μL RNaseOUT, 0.5 μL Superscript III (Invitrogen)). Reactions were incubated 5 minutes at 25 ºC, 20 minutes at 42 ºC, 40 minutes at 50 ºC, 5 minutes 80 ºC. After retro-transcription, the samples were loaded in a 15% denaturing polyacrilamide-urea gel and size selected. Next, the cDNA was extracted from the gel and precipitated.

### Recircularization and PCR amplification

cDNA was resuspended in 15 μL of water, 2 μL CircLigase Reaction buffer 10x, 1 μL ATP, 1 μL MnCl^2^ 50 Mm and 1 μL of Circligase ssDNA ligase (Epicentre). Reactions were incubated 2 hours at 60 ºC. After recircularization, the libraries were amplified using primers compatible with Illumina adaptors (Forward oligo AATGATACGGCGACCACCGAGATCTACAC, reverse oligo CAAGCAGAAGACGGCATACGAGATcgtgatGTGACTGGAGTTCAGACGT-

GTGCTCTTCCGATCT, where lower case represents one of the barcodes) and the PCRs products were purified in a 10% non-denaturing polyacrilamide gel. Libraries were sequenced at the Yale Center for Genome Analysis using HiSeq2500 platform. Raw reads are publicly accessible in the Sequence Read Archive under SRPxxx.

### Label-transfer assay

Zebrafish embryos were injected at one-cell stage with *in vitro* transcribed mRNA encoding for the indicated RBPs. Embryos where collected at 4 hpf and processed following the same procedure described for iCLIP samples, including UV-cross linking, RNA digestion, FLAG pull-down, radiolabeling, protein gel and transfer, with the only modification that in this case a higher concentration of RNAseI (Invitrogen) was used (1:500 final dilution). After protein transfer, membranes were exposed to a phosphorimager storage screen to detect the radioactive signal from the RNA. After exposure, Western blot was performed on the membranes to detect FLAG-tagged RBPs. Rabbit anti-FLAG antibody (Sigma) was used as a primary antibody in the Western blot.

### Khsrp iCLIP with endogenous antibody

iCLIP experiment as described in Huppertz et al, 2014 with minor changes. Embryos at sphere stage (4 hpf) were collected and irradiated with 254 nm UV light to induce crosslinking (embryos were snap frozen and stored in batches to yield a total of 1,000 embryos/condition). Frozen embryos were then thawed and homogenized on ice in iCLIP lysis buffer. An affinity-purified rabbit polyclonal antibody raised against zebrafish KHSRP (generated by YenZym Antibodies, LLC) was used to isolate RNA-protein complexes. Briefly, 200 µL Protein G Dynabeads were added to 50 µg antibody in lysis buffer. Beads were incubated for an hour, washed three times with lysis buffer and added to the lysates. As a control, a parallel experiment was performed without the addition of antibody. Subsequent steps in the iCLIP protocol were performed as described above. Twenty cycles of amplification of cDNA were used for final library construction. Barcoded PCR-amplified libraries were size-selected in a 6% TBE gel and combined for Illumina sequencing. Both the KHSRP pull-down and the no-antibody control were performed in triplicates.

### iCLIP analysis: Binding profiles

#### Demultiplexing samples

After sequencing of the iCLIP libairies, raw reads were composed as NNNN-insert-NN-barcode(4-mer)-adapter where (i) the 6N (NNNN+NN) are random nucleotides composing the Unique Molecular Identifier (UMI), (ii) the 4mer barcode is the in-house barcode used to demultiplex samples and (iii) adapter is the 3’ Illumina adapter. To demultiplex samples barcoded with in-house barcode, reads were first trimmed of Illumina adapter by aligning its sequence, requiring 100% match of the first five base pairs and a minimum global alignment score of 80 (Matches: 5, Mismatches: −4, Gap opening: −7, Gap extension: −7, Cost-free ends gaps). Only trimmed reads were then demultiplexed, allowing a single mismatch with the expected barcodes. The 5’ 4N and 3’ 2N composing the UMI were then clipped and added to the read name for downstream analysis. Reads with UMI containing unidentified nucleotide(s) were discarded.

#### Mapping

Demultiplexed reads were mapped to the zebrafish Zv9 genomes using STAR (Dobin et al, 2013) version 2.4.2a with the following non-default parameters: *╌alignEndsType EndToEnd ╌outFilterMultimapNmax 100 ╌seedSearchStartLmax 15 ╌sjdbScore 2*. Genomic sequence indices for STAR were built including exon-junction coordinates from Ensembl r78 (Aken et al, 2017).

#### UMI

To filter unique molecules, reads were considered unique when they had a unique UMI, as well as unique start and end positions.

#### Binding profiles

Using Ensembl 78, per chromosome and per transcript annotation binding profiles were created by summing the first position (5’ end) of reads.

### RBP binding metagene profiles

To summarize RBP binding within protein-coding transcripts, all UTRs and CDSs iCLIP profiles were split into 50 bins (i.e. bins are not equally sized). Transcripts with less than 50 reads total or less than 10 reads within a UTR or CDS were discarded. Bins were then summed from the most 5’ to the 3’ end for all transcripts (first bin of first transcript with first bin of second transcript, etc.) to obtain metagene profiles. Hierarchical clustering with complete-linkage method using an Euclidean distance matrix was applied to group similar binding profiles together. For clustering, biological replicates were merged together by summing replicates.

### RBP motif and overlap heatmap

To define binding preference of RBP, bound positions were first defined within transcripts for nucleotide with at least 2 unique reads. These positions extended 5 nucleotides upstream and 10 nucleotides downstream defined bound windows. In each window, 6mers were counted. Transcriptome-wide 6mer frequencies were then normalized per RBP. To take into account the iCLIP experimental background, these frequencies were normalized by the average 6mer frequencies measured for the 6 controls. For the khsrp iCLIP experiment using an endogenous antibody, a minimum of 10 unique reads was required to define bound windows to take into account the higher sequencing depth of these samples, while the cutoff for the controls remained at 2. To build the logo representation, the top 10 most frequent normalized 6mer were aligned with MAFFT (Katoh et al, 2013) using the parameters *╌reorder ╌lop -10 ╌lexp -10 ╌localpair ╌genafpair ╌maxiterate 1000*. The multiple sequence alignment positions with more than 50% gaps were trimmed on both ends. Nucleotide frequencies were calculated using the final trimmed alignment, and represented as logo. To cluster RBP based on logo, the overlap of the top 20 normalized 6mer frequencies with iCLIP controls were used to perform hierarchical clustering similarly to metagene profiles. Biological replicates were also merged together by summing replicates.

### Motif-centered metaplots

To simultaneously analyze binding and regulatory activity, sequences matching RBP motifs were searched within transcriptome. Windows were defined 100 nt upstream and downstream of the motif occurrence. Windows with minimum RESA coverage below 0.05 CPM in one of the 2 conditions used (e.g. 6 hpf and 2 hpf) were discarded. Finally, iCLIP (5’end) and RESA (coverage ratio) windows were averaged. To define RBP binding context, all 3-mer were counted with 20 nt window upstream and downstream of the motif (not including it). The 10% most regulated motif occurrences were used to define the 3-mer frequencies of favorable context, while the 10% least regulated defined unfavorable context. Strong sites had a favorable context and no unfavorable context, while weak or control sites had neither. For each set of sites, the corresponding iCLIP signal was averaged before plotting.

### Computational model training

The RESA profiles were analyzed using a sliding window approach: a window of 100 nucleotides (nt) was used with a sliding step-size of 10 nt to scan the profile of each transcript. For each window, the frequency of all kmers was computed (1-8 nt). The RESA stability value of the center of each window (the 50^th^ nt) was computed, which was used to represent the stability of that window. After scanning all the profiles, this window-based data was used to train a Random Forest (RF) model with 500 trees (R 3.4.2, package of “*randomForest*” 4.6-12), where the objective of the training process is to build a set of decision trees that are able to capture the association between kmer frequencies and their correlation with RESA stability values. The trained RF model was used to predict the stability of new transcripts using only their sequence data in a similar window scanning procedure: For each new transcript, a scanning window of 100 nt and slide size 10 nt was used, where the trained RF model was used to predict the stability of each window using the kmer frequencies data computed per window. The predicted RESA values of all the sliding windows spanning the whole profile constitute the predicted RESA profile for this transcript.

### Feature Selection

In order to make the learning process computationally feasible, a pre-processing kmers filtering step was required to decrease the large number of kmers (87,380 kmers) while maintaining the potentially important ones as follows: First, all the kmers with non-zero occurrence frequency were retained (31,548 kmers). Second, the kmers with absolute correlation (Spearman correlation coefficient) greater or equal to 0.1 (with respect to RESA stability values) were selected as potentially correlated (387 kmers). Finally, of these pre-selected kmers, the trained RF model highlighted that 57 motifs had the most significant contribution.

### 5-fold Cross validation

To assess how the trained model would generalize to an independent dataset, a 5-fold cross validation was used to evaluate the predictive model performance in practice. All the transcripts were randomly partitioned into 5 equal sized subsamples. Of these 5 subsamples, a single subsample was retained as the validation set of transcripts for testing the RF model, while the remaining 4 subsamples were used as training data for building the RF model. This process of training and validation was then repeated 5 times, with each of the 5 subsamples used exactly once as the validation data. Model performance assessment through cross-validation is performed for each library (UTR targeted and poly(A)-selected), separately. For each of these two libraries, a separate RF model was trained and validated for each different treatment conditions (WT, LNA-430, and *α*-amanitin), individually.

### Selected motifs analysis

For each selected motif, the differential stability between windows that contain that motif and those that don’t was compared. The mean RESA values between these two groups of windows (motif presence vs. absence) was used as a proxy to reflect the average differential effect of each individual motif on stability. All the windows were grouped into three categories according to their RESA stability profile; the top 25% (Stable), middle 50%, and bottom 25% (Unstable). For each selected motif, the motif mean frequency of each group (top 25% Stable and bottom 25% Unstable) was compared against the middle group, which highlighted the preferential enrichment of each motif in the stabilizing/destabilizing groups compared to the middle, control group.

### Model performance on endogenous transcripts

To evaluate the RF model performance in predicting stability of endogenous mRNAs, the model that was trained on average RNA-seq stability profiles from the RESA library was used. The motifs frequencies across the 3’-UTR of each endogenous transcript was used as input to the trained model. Based on motifs frequencies data, the RF model predicted the fold-change of each endogenous mRNA, which was compared against the fold-changes measured with mRNA-seq.

### Model validation on trip10 locus

#### Model training

To avoid any bias in the model training, the RF model was trained on all the RESA targeted transcripts after excluding the *trip10* transcript. After training, the model performance was assessed by comparing the experimental profile of *trip10* transcript according to RESA against the predicted profile according to the trained RF model.

#### Reporter validation trip10

WT

NNNNAAATAATATAATTTATTGAGTAAATaagcgcttGTATATTAAATAAACATGTATGTAAGANNNN

WT_F

ccctacacgacgctcttccgatctNNNNAAATAATATAATTTATTGAGTAAATaagcgcttGTATA

WT_R

gAGTTCAGacgtgtgctcttccgatctNNNNTCTTACATACATGTTTATTTAATATACaagcgcttATTTACTCAAT

Mut1

NNNNAAATAATATAATTTATTGAGTAAATaaAAgGttGTATATTAAATAAACATGTATGTAAGANNNN

Mut1_F

ccctacacgacgctcttccgatctNNNNAAATAATATAATTTATTGAGTAAATaaAAgGttGTATA

Mut1_R

gAGTTCAGacgtgtgctcttccgatctNNNNTCTTACATACATGTTTATTTAATATACaaCcTTttATTTACTCAAT

Mut2

NNNNAcATAAcATcActcATcGAGTAAATaagcgcttGTATAccAcATAcACATGcATGTAAGANNNN

Mut2_F

ccctacacgacgctcttccgatctNNNNAcATAAcATcActcATcGAGTAAATaagcgcttGTATA

Mut2_R

gAGTTCAGacgtgtgctcttccgatctNNNNTCTTACATgCATGTgTATgTggTATACaagcgcttATTTACTCgAT

Mut3

NNNNAcATAAcATcActcATcGAGTAAATaaAAgGttGTATAccAcATAcACATGcATGTAAGANNNN

Mut3_F

ccctacacgacgctcttccgatctNNNNAcATAAcATcActcATcGAGTAAATaaAAgGttGTATA

Mut3_R

gAGTTCAGacgtgtgctcttccgatctNNNNTCTTACATgCATGTgTATgTggTATACaaCcTTttATTTACTCgAT

Mut4

NNNNAAgcAAgcTAgcTgcTTGAGgcAAgcaAAgGttGgcTATgcAAgcAACATGgcTGgcAGANNNN

Mut4_F

ccctacacgacgctcttccgatctNNNNAAgcAAgcTAgcTgcTTGAGgcAAgcaAAgGttGgcTA

Mut4_R

gAGTTCAGacgtgtgctcttccgatctNNNNTCTgcCAgcCATGTTgcTTgcATAgcCaaCcTTtgcTTgcCTCAAg

sticky ends_F 10353

CCTGACTCGACTCGAGCCCTACACGACGCTCTTC

sticky ends_R 10354

TCACTATAGTTCTAGAGAGTTCAGACGTGTGCTC

Hi-Fidelity PCR (using Phusion NEB-F-530S) was run with Forward and Reverse oligos. The PCR product was run on gel then extracted and purified using Qiagen Gel Extraction Kit. Another PCR was run to add the “sticky ends” to each of the previous PCR products. The final product was cloned into pCS2-GFP vector (cut with XhoI-XbaI) using In-Fusion^®^ HD Cloning Plus-Clonetech-639642). Each construct was sequenced to confirm the cloning of the expected sequence. Each of the final *trip10* plasmids was then linearized with NotI and used as a template for *in vitro* transcription using (SP6 mMessage Machine-Ambion AM1340). To test the effect of each trip10 sequences on gene expression, a mix of 100 pg trip10 and 70 pg DsRed mRNA as a control was injected for each trip10 sequence into one-cell stage embryos as described previously. The injected embryos were incubated for 24h at 28 °C, and the GFP expression was compared to the control DsRed expression using fluorescence microscopy. To measure the effect of each trip10 sequence on mRNA stability, a second set of embryos was injected at the one-cell stage with an mRNA cocktail containing 5 pg of each of the GFP trip10 sequences. Then, 30 embryos were collected at 64-cell and shield stages (3 replicates each). Frozen embryos were lysed with 1 mL of Trizol and total RNA was extracted following the manufacturer’s protocol. totRNA was dissolved in 11 µL of water and reverse transcribed using SuperScript III with 100 nM of reverse transcription primer (5’-CTTCAGACGTGTGCTCTTCCGATCTNNNNNNNNNN-barcode-CATGTCTGGATCTACGTAATACG-3’) in 20 µL reactions following the manufacturer’s protocol. 64-cell and shield samples were reverse transcribed with primers containing the 5’-TAGCTT-3’ and 5’-ACTTGA-3’ homemade barcodes, respectively. After reverse transcription, 64-cell and shield samples were pooled per-replicate and cDNAs were purified using 108 µL of AMPure XP beads (Beckman Coulter Inc.) following the manufacturer’s protocol and, then, dissolved in 20 µL of water. 5’- and 3’-Illumina adapters were added by PCR with the follow primers (forward 5’-AATGATACGGCGACCACCGAGATCTACACTCTTTCCCTACACGACGCTCTTCC-3’ and reverse: 5’-CAAGCAGAAGACGGCATACGAGAT-illumina_barcode-GTGACTGGAGTTCAGACGTGTGCTCTTCC-GATCT-3’) and amplified using the lowest amount of PCR circles necessary to obtain enough material for sequencing. Replicate 1, 2 and 3 were amplified using the 5’-CCACTC-3’, 5’-ATCAGT-3’ and 5’-AGGAAT-3’ Illumina barcodes, respectively. PCR products were precipitated using ethanol and purified in an 8% non-denaturing polyacrylamide gel. Libraries were sequenced at the Yale Center for Genome Analysis using HiSeq2500 platform. Raw reads are publicly accessible in the Sequence Read Archive under SRPxxx.

#### Demultiplexing samples

The sequenced pairs had the following structure: R1-4N-insert-4N-plasmid-barcode(6-mer)-10N-R2. The second read R2 was used to demultiplex the samples while R1 was used to measure the WT and mutant sequences. R2 was composed as NNNNNNNNNN-barcode(6-mer)-plasmid where (i) the 10N are random nucleotides composing the Unique Molecular Identifier (UMI), (ii) the 6mer barcode is the in-house barcode used to demultiplex samples. To demultiplex samples barcoded with in-house barcode, reads were first clipped of 10N on their 5’ end composing the UMI and added to the read name for downstream analysis. Reads with UMI containing unidentified nucleotide(s) were discarded. Reads were then demultiplexed, allowing a single mismatch with the expected barcodes.

### Mapping

The first read of demultiplexed read-pair was composed as NNNN-insert-NNNN-plasmid where the 4N are random nucleotides. First reads were (i) clipped of 4N on their 5’ end, (ii) trimmed on their 3’ end of the sequence AGATCGGAAGAGCACACGTCTGAACTCTCTAGA by alignment, requiring 100% match of the first five base pairs and a minimum global alignment score of 60 (Matches: 5, Mismatches: −4, Gap opening: −7, Gap extension: −7, Cost-free ends gaps), (iii) clipped of 4N on their 3’ end, then (iv) mapped to the *trip10* WT and mutant sequences using Bowtie2 2.3.4.1 (Langmead et al, 2012) with the following non-default parameter: *╌score-min L,-0.6,-0.1*. Because WT and mutant sequences were similar, this parameter limited the number of allowed mismatch to get unique mapping of first reads on WT and mutant sequences.

### UMI counts

To filter unique molecules, reads were considered unique when they had a unique UMI. Reads with unique UMI were counted for WT and mutant sequences. Counts were then normalized to CPM (Count Per Million) using total counts of all WT and mutant sequences per sample.

---

**Figure S1.**
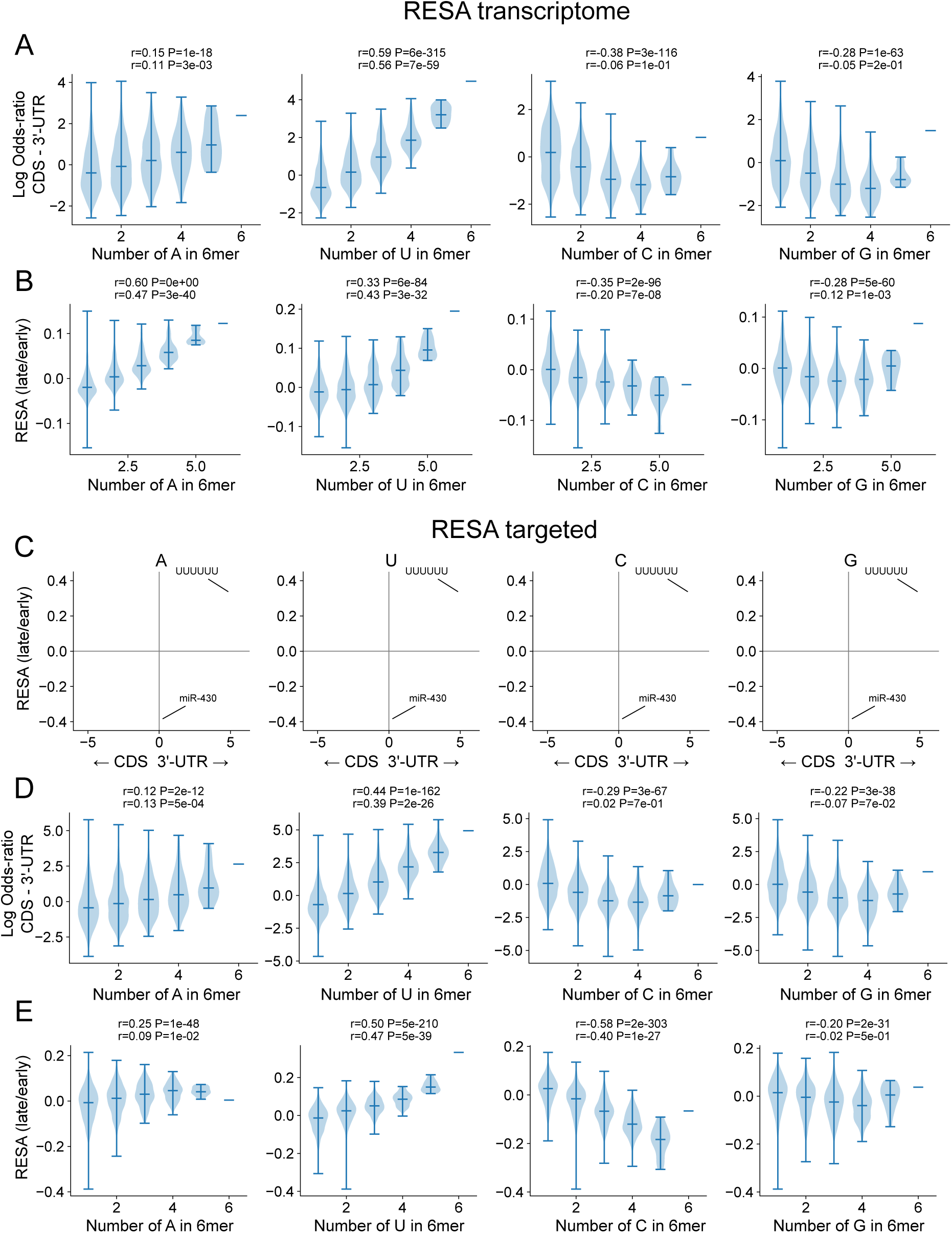
(A, D) (A, D) Violinplot comparing the number of A, U, C and G nucleotides in all 6mers and their log odds-ratio within CDS vs. 3’-UTR for RESA transcriptome (A) and targeted (D). Positive log odds-ratio corresponds to enrichment in 3’-UTR. Pearson correlations are indicated for 1 to 6 nucleotides (top) and 3 to 6 nucleotide (bottom) per 6mer. (B, E) Violinplot comparing the number of A, U, C and G nucleotides in all 6mers and their RESA average stability between 2hpf and 6hpf for RESA transcriptome (B) and targeted (E). Same correlations as (A, D). (C) Biplots comparing all hexamers relative enrichment within CDS and 3’-UTR to their average coverage ratio between 2 hpf and 6 hpf for RESA targeted library. From left to right, biplot is colored with number of U, A, C and G within each hexamer.

**Figure S2.**
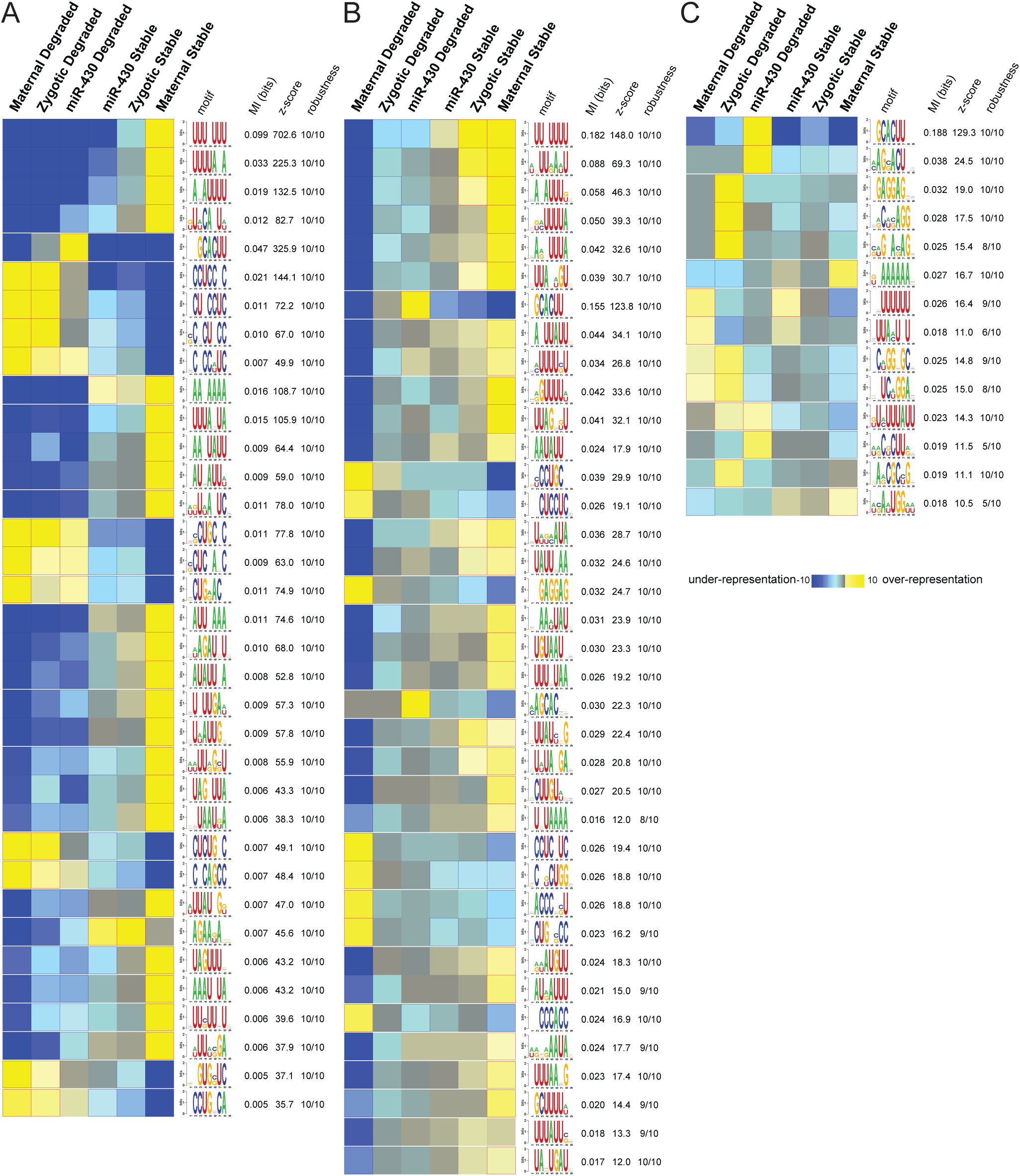
Full list of linear motifs discovered by FIRE that were informative of the various modes of regulation across the RESA libraries (A) for the transcriptomic library, (B) the targeted library and (C) the poly(A) selected targeted library. Shown are each motif’s id, its primary sequence in a weblogo format, its associated mutual information value and Z-score (estimated using a randomization-based statistical test; Elemento et al, 2007), and its robustness score from a three-fold jackknifing test (Elemento et al, 2007). Yellow entries denote enrichment while blue entries mark significant depletion of a given motif in each corresponding cluster. Z-score cut-off of 20 for the transcriptomic and 10 for the targeted libraries.

**Figure S3.**
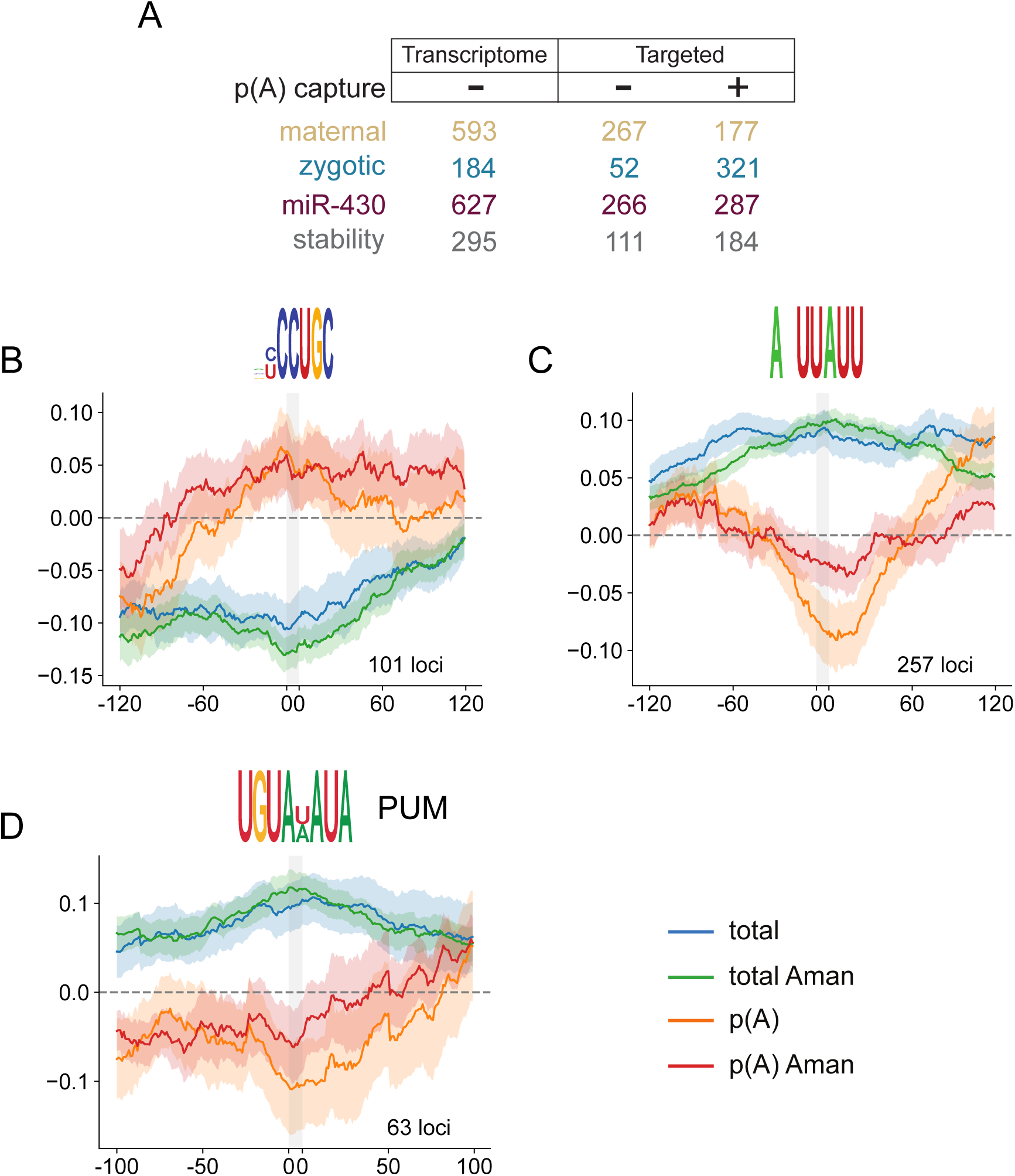
(A) Sliding windows across transcriptome-based library or high-coverage targeted library (with/without poly(A) selection) identify sequences under different modes of regulation. (B-D) Motif-centered metaplots for CCUGC (B), ANUUAUU (C) and Pumilio (D) motifs. Targeted RESA libraries coverage ratio were averaged over windows centered on RBP motif (RESA minimum coverage >0.01 CPM). Motif is represented with grey bar. S.E.M. of RESA is shown by shaded outlines.

**Figure S4.**
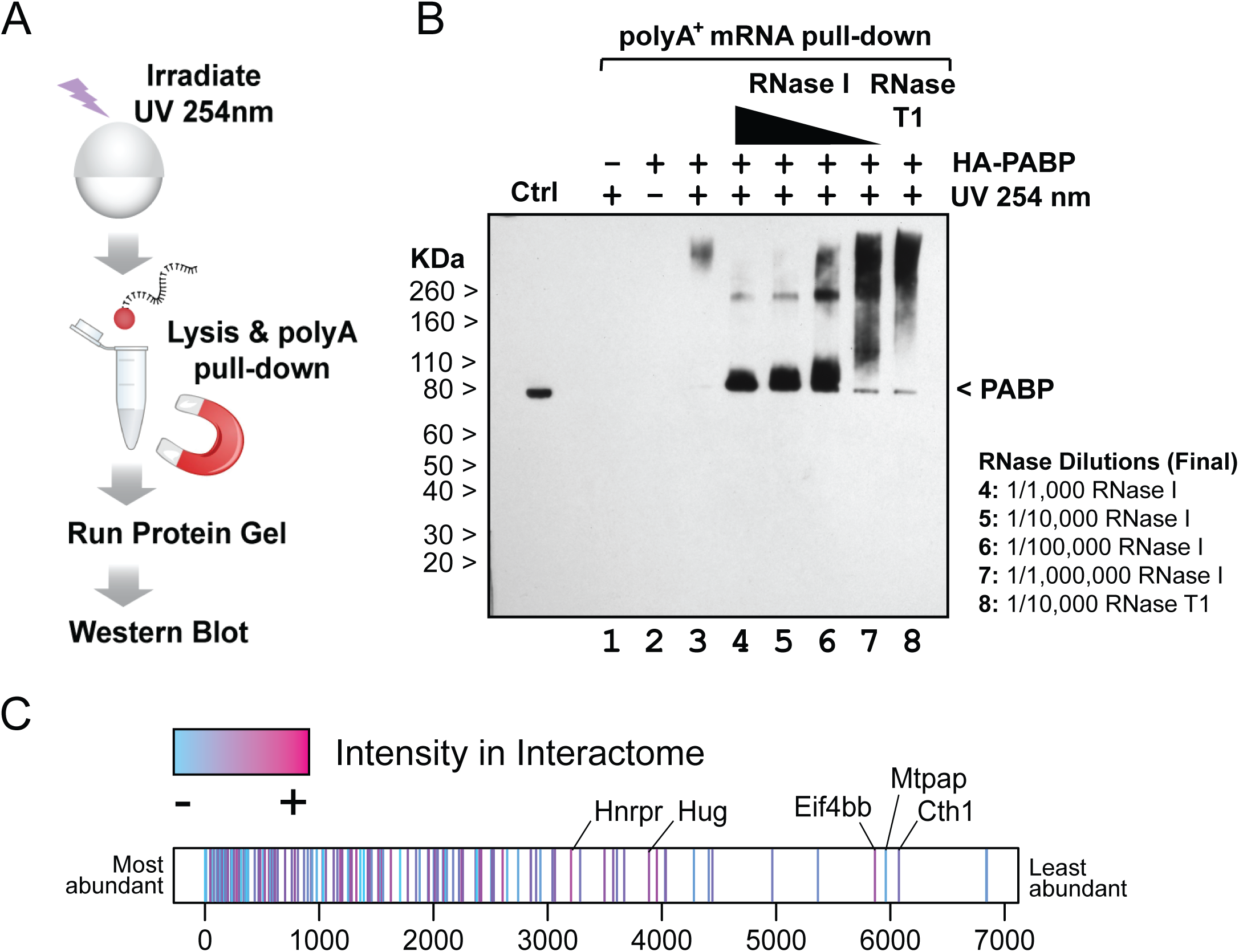
Schematic presentation of the experimental procedure for interactome capture (B) Efficiency/specificity of capture after *in vivo* UV crosslinking in embryos was assessed by measuring capture efficiency of an HA-tagged variant of PABP, an abundant protein that binds to the 3’ ends of poly(A) transcripts. (C) Interactome capture identifies RBPs from all the range of expression levels. Proteins identified in the input were ranked according to their abundance in the whole-embryo lysate. Proteins identified in the interactome capture were then labeled according to their intensity in the interactome capture. Although overall abundance of individual proteins (input) generally correlated with capture efficiency, several of the most highly captured proteins were lowly expressed.

**Figure S5.**
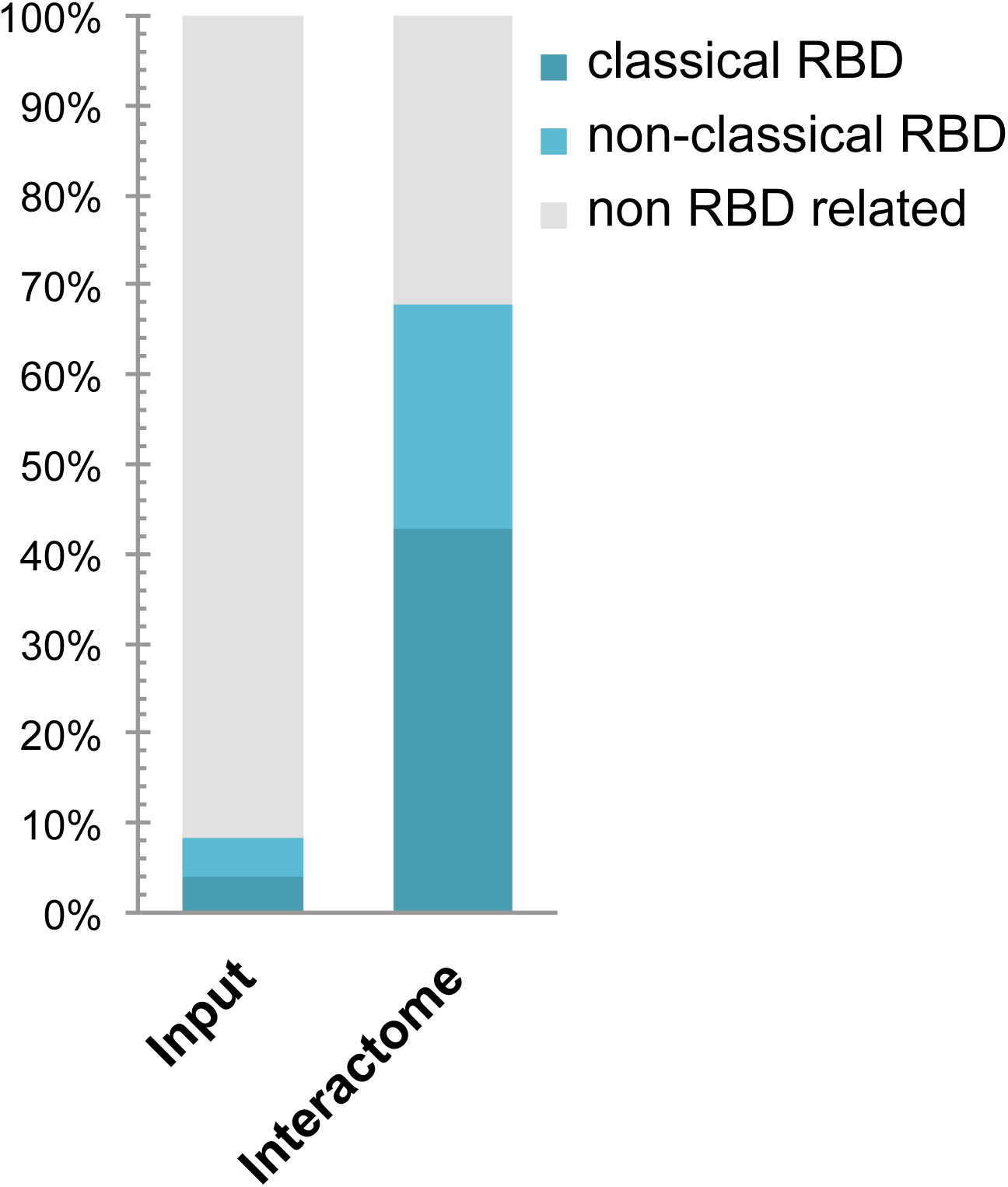
Enrichment of proteins containing RNA-binding domains (RBD) in interactome capture. Proteins identified in the input and in the interactome capture were classified in three categories regarding the annotation of their Interpro domains as RBDs in Castello et al, 2012. If a protein contains a mixture of classical and non-classical domains, it is only counted once and only in the classical RBD category.

**Figure S6.**
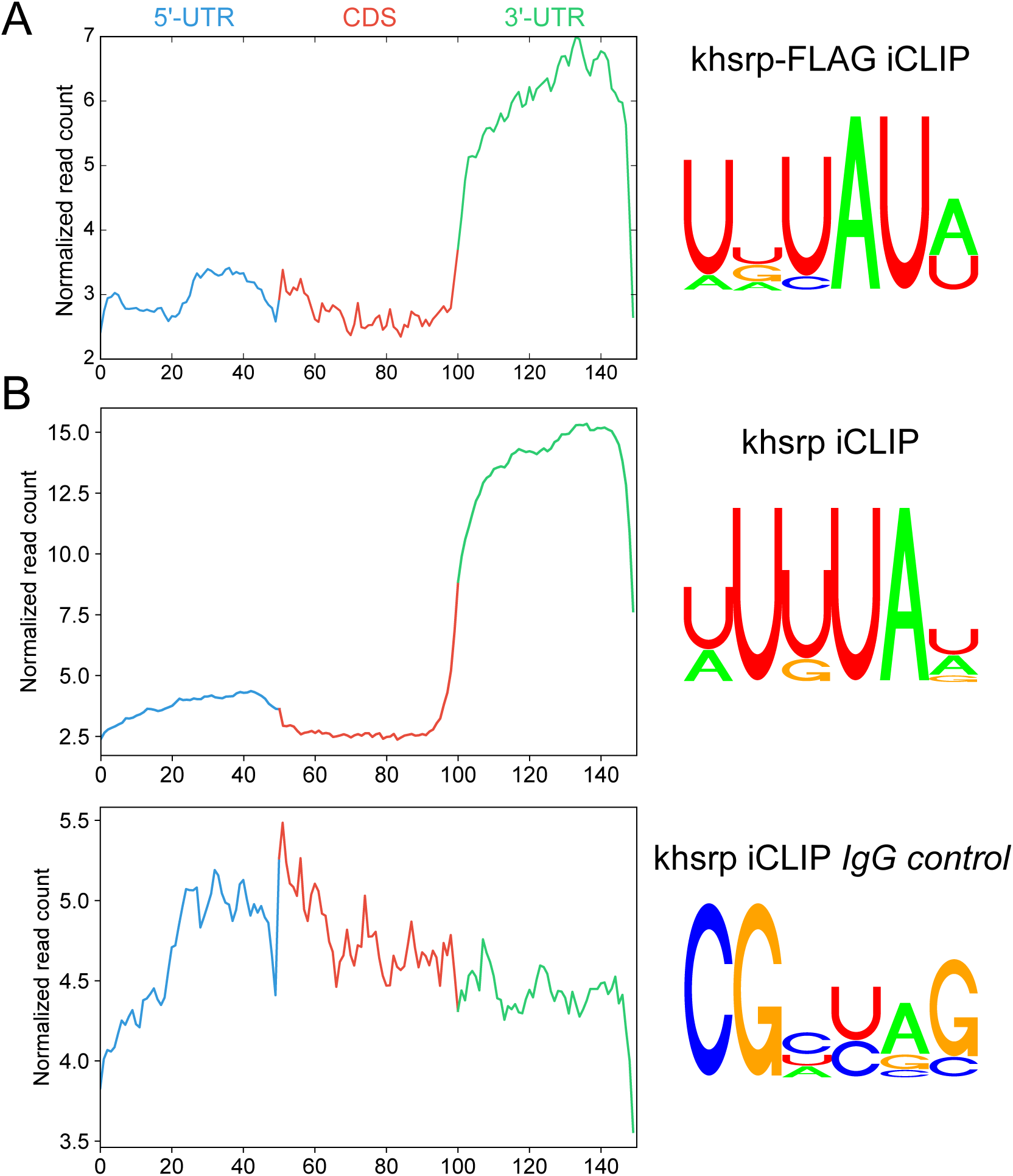
Comparison of *khsrp* iCLIP with FLAG-tag (A) and endogenous (B) proteins. iCLIP metaplots of RBP-binding within protein-coding transcripts (left). Weblogo representation of the top 6-mer normalized by respective iCLIP control (right).

**Figure S7.**
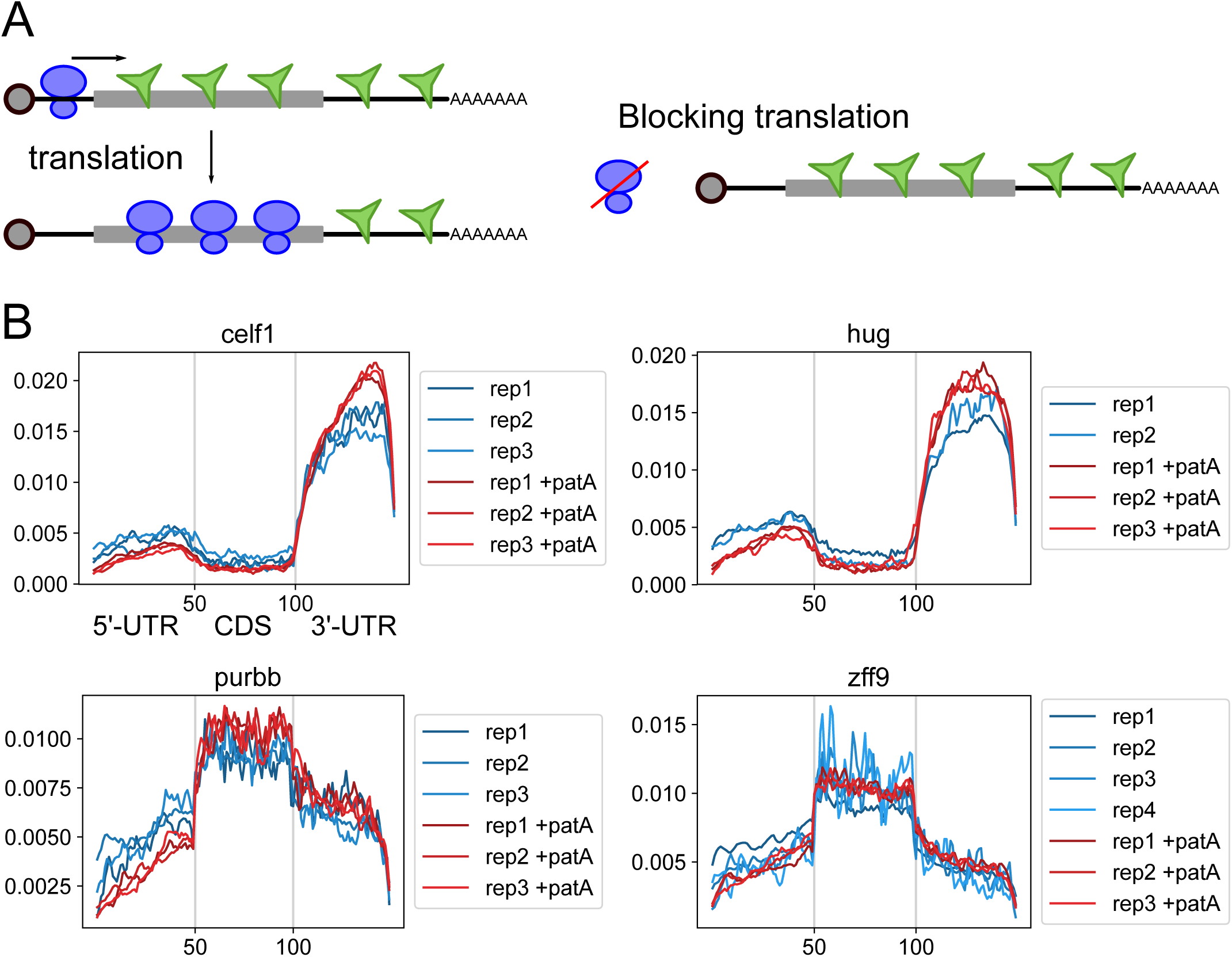
Testing translation influence on RBP binding. The differential binding between coding and non-coding regions of RBPs could be induced by the translating ribosome preventing stable binding to the coding regions. Alternatively, the binding pattern might be determined by the preferential accumulation of binding motifs within those regions. To distinguish between these scenarios, translation was inhibited using pateamine A, which prevents the formation of the translational initiation complex by inhibiting *eif4A*. Comparison of transcriptome metaplots for *celf1*, *hug*, *purbb* and *zff9* in replicates in WT and patA treated embryos revealed similar enrichment in 3’-UTR regions after translation inhibition. (A) Diagram of hypothesis tested. Translating ribosomes would actively push RBPs out of the coding-sequence that would be blocked in presence of translation inhibitor patA. (B) iCLIP metaplots of RBP-binding within protein-coding transcripts comparing WT and patA treated embryos for *celf1*, *hug*, *purbb* and *zff9* in replicates.

**Figure S8.**
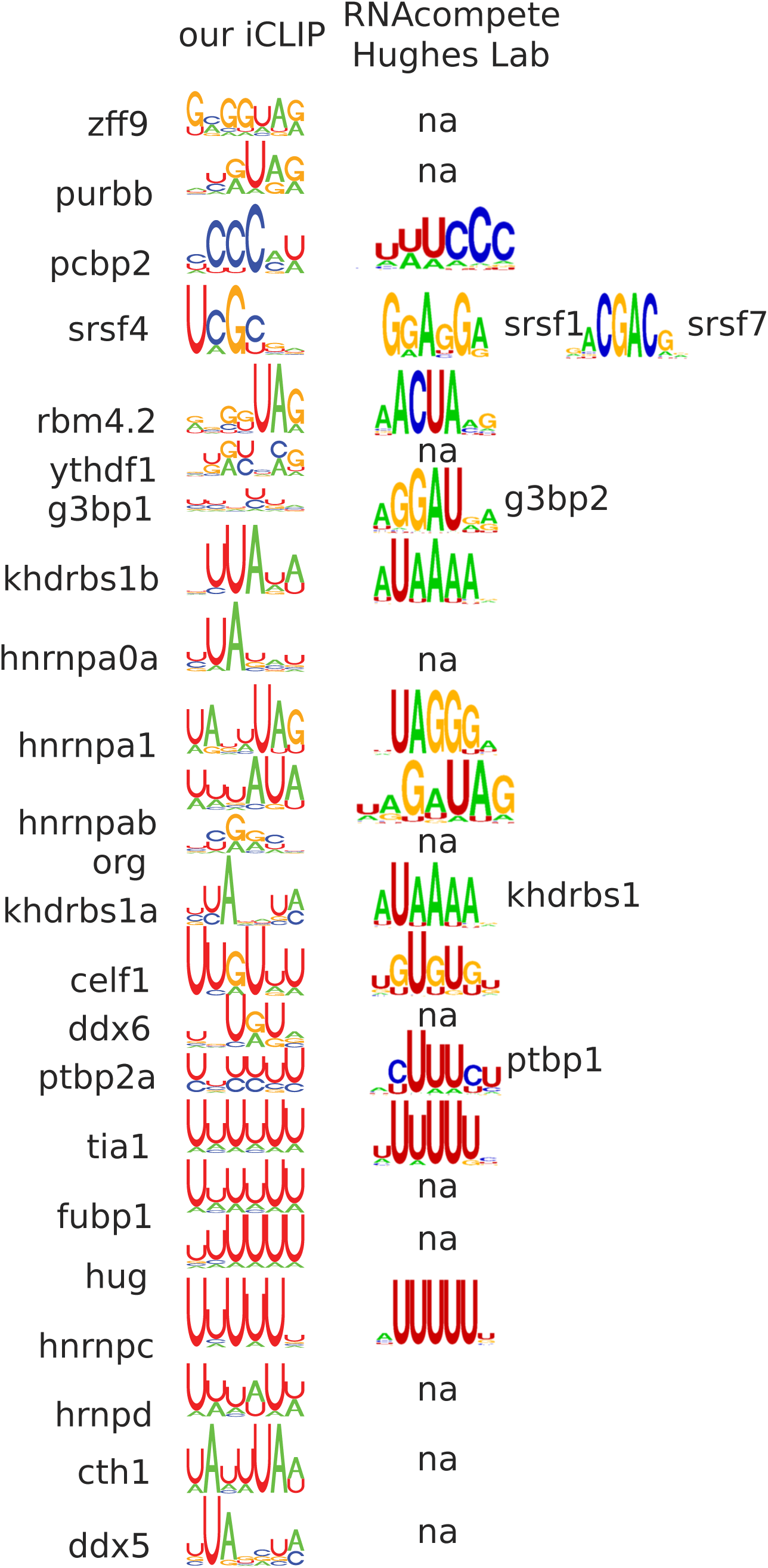
Comparison of our iCLIP-derived motif with Ray et al, 2013 assay. When only a related protein was available, the name is shown. (na: not-available protein in Ray et al, 2013 database).

**Figure S9.**
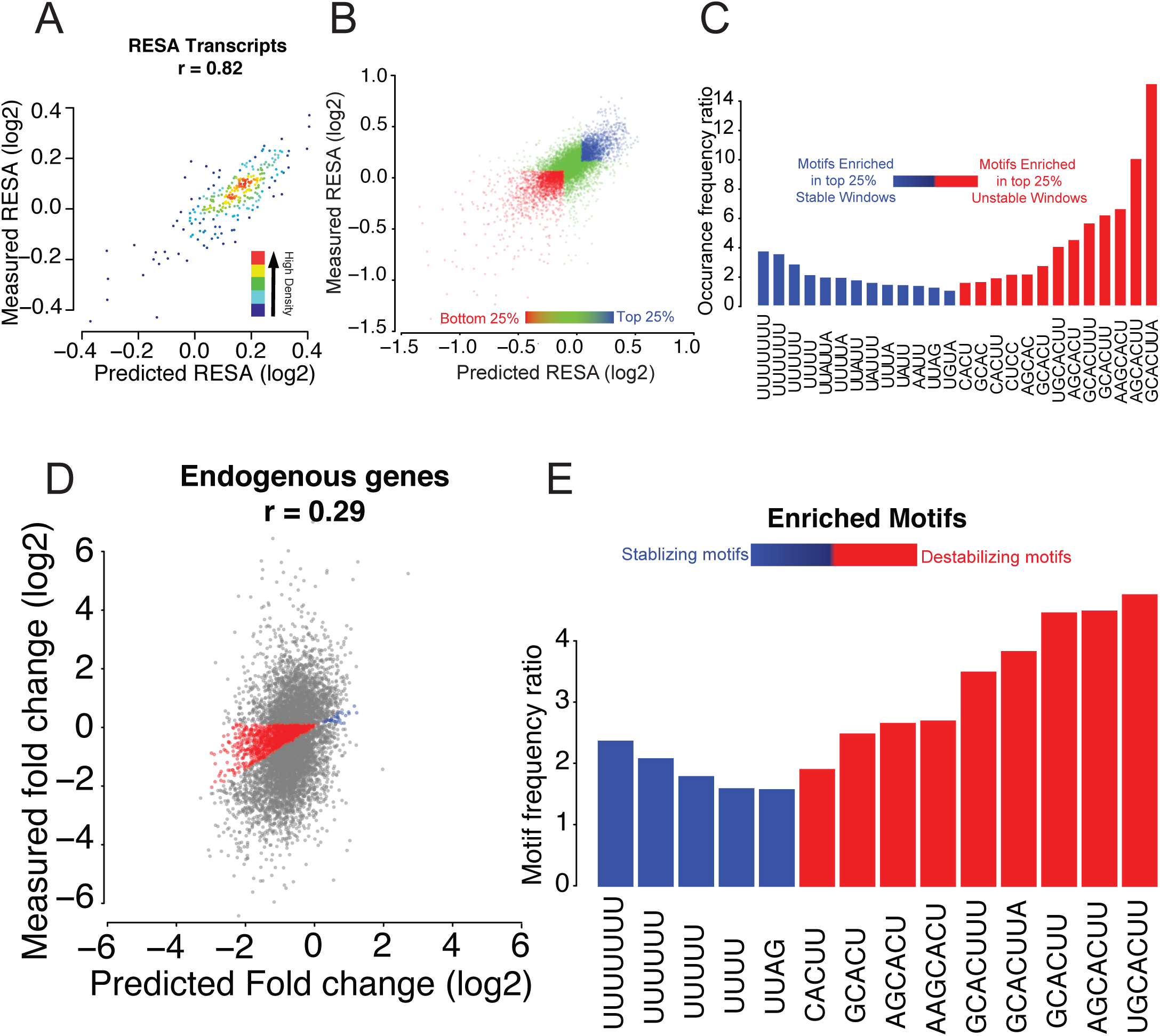
RESA Random forest model. (A) Model performance per transcript using 5-fold cross validation. Model achieved a 0.82 Pearson correlation between mean predicted stability of sliding windows per transcript and mean measured stability according to RESA profile. Colors represent density (red, yellow, green, light blue, dark blue, low density). (B) Comparing mean motifs frequencies ratios in top 25% stabilizing and top 25% destabilizing windows (blue, and red dots, respectively) against the middle control 50% windows (green dots). (C) Motifs enriched in the top 25% stabilizing and top 25% destabilizing windows (blue and red bars, respectively). Y-axis represents per motif occurrence frequency ratio between top 25% stabilizing/destabilizing windows and middle 50% windows. All the motifs selected have statistical significant *P*-values below 2.2e-18 (Mann-Whitney U-test followed by Bonferroni multiple testing adjustment). (D) Model performance on endogenous transcripts. Model achieved a 0.29 Pearson correlation between predicted fold change and measured fold change according to mRNA-seq. 18% of the transcripts (red and blue dot for destabilizing and stabilizing transcripts, respectively) were accurately predicted (same fold-change within 10% error range). (E) Motifs enriched in the 18% accurately predicted endogenous transcripts using the computational model trained on RESA library. Y-axis represents the occurrence frequency ratio between the accurately predicted transcripts (red and blue dots) against the less accurately predicted transcripts (gray dots). All the motifs selected have statistical significant *P*-values below 0.05 (Mann-Whitney U-test followed by Bonferroni multiple testing adjustment).

**Figure S10.**
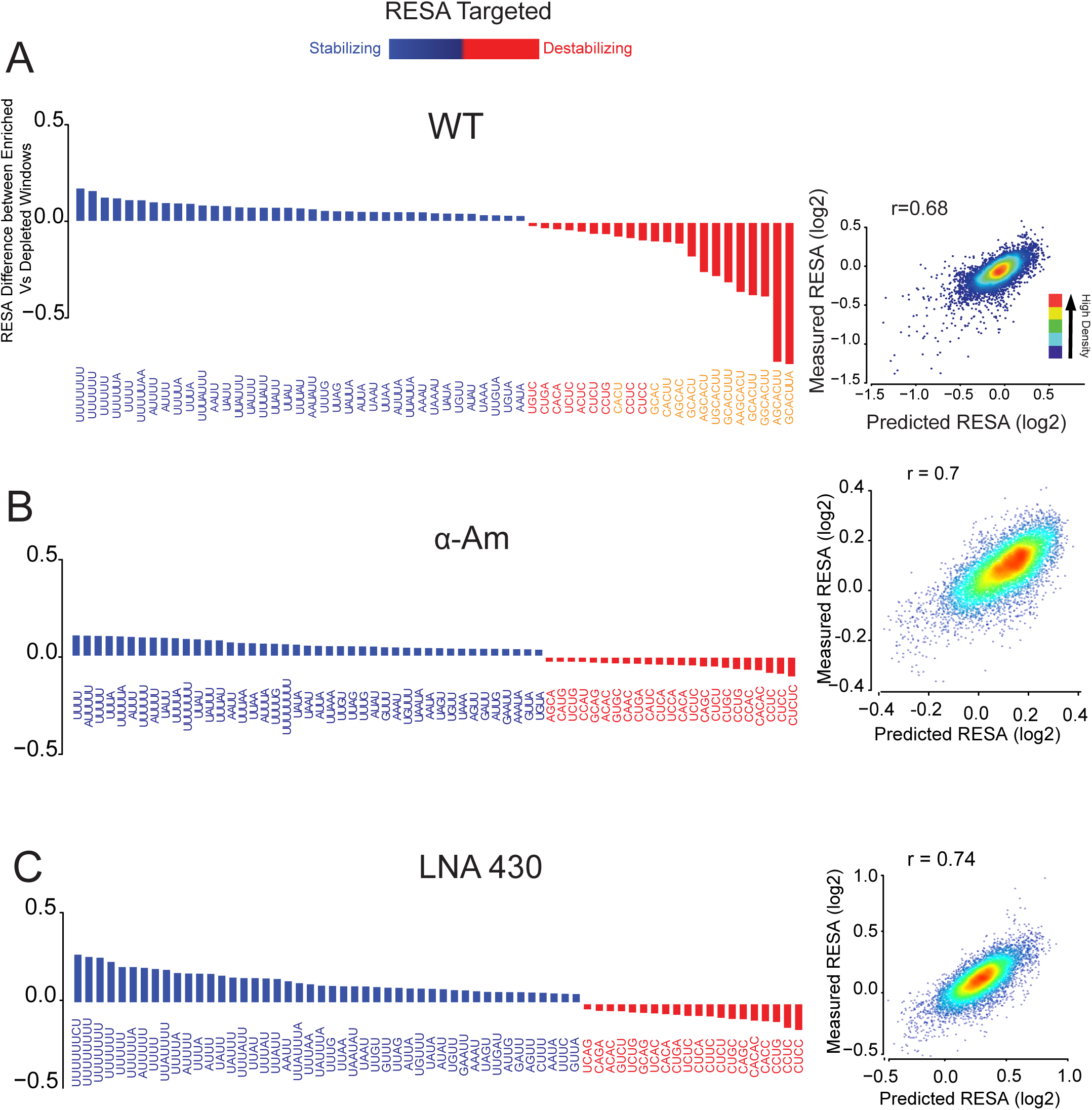
Random forest based motif selection for RESA Targeted library. (A) (left) Top selected motifs according to the trained model on WT mRNAs. Y-axis represents the RESA fold change difference between windows that contain each motif and those that don’t. All the motifs selected have statistical significant *P*-values below 4e-42 (Mann-Whitney U-test followed by Bonferroni multiple testing adjustment). (right) Model performance per window using 5-fold cross validation; Model achieved a 0.68 Pearson correlation between predicted stability (x-axis) and measured stability (y-axis) according to RESA (Colors represent density (red, yellow, green, light blue, dark blue, low density)). (B) Same as (A), but the model was trained and tested after blocking transcription with RNA polII inhibitor, *α*-amanitin. (C) Same as (A), but the model was trained and tested after miR-430 was inhibited using an antisense tiny-LNA complementary to miR-430 (LNA^430^).

**Figure S11.**
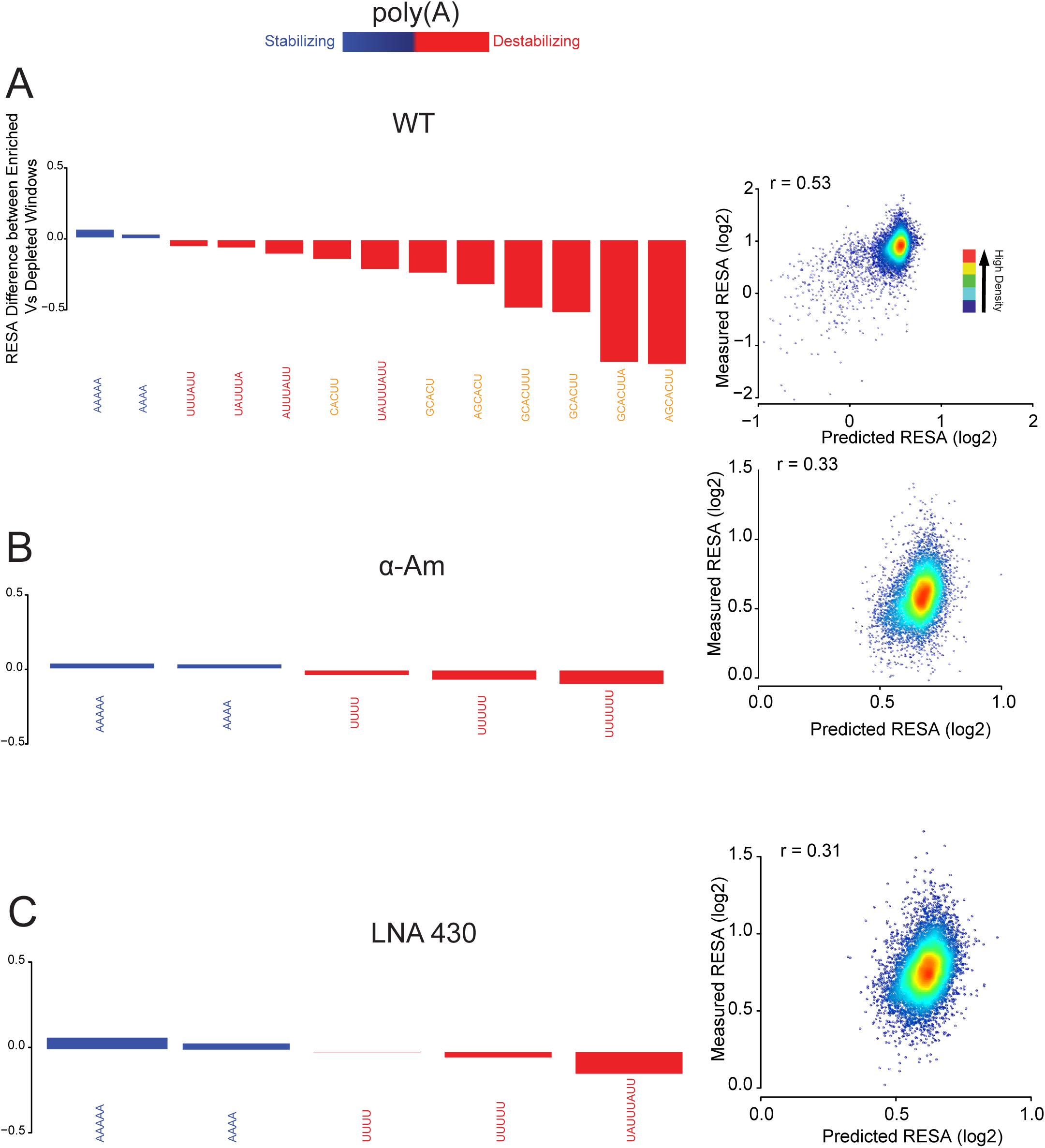
Random forest based motif selection for RESA poly(A) selected library. (A) (left) Top selected motifs according to the trained model. Y-axis represents deadenylation difference between windows that contain each motif and those that don’t. All the motifs selected have statistical significant *P*-values below 0.01 (Mann-Whitney U-test followed by Bonferroni multiple testing adjustment). (right) Model performance per window using 5-fold cross validation; Model achieved a 0.53 Pearson correlation between predicted deadenylation (x-axis) and measured deadenylation according to RESA library (y-axis). Colors represent density (red, yellow, green, light blue, dark blue, low density). (B) Same as (A), but the model was trained and tested after blocking transcription with RNA polII inhibitor, *α*-amanitin. (C) Same as (A), but the model was trained and tested after miR-430 was inhibited using an antisense tiny-LNA complementary to miR-430 (LNA^430^).

**Figure S12.**
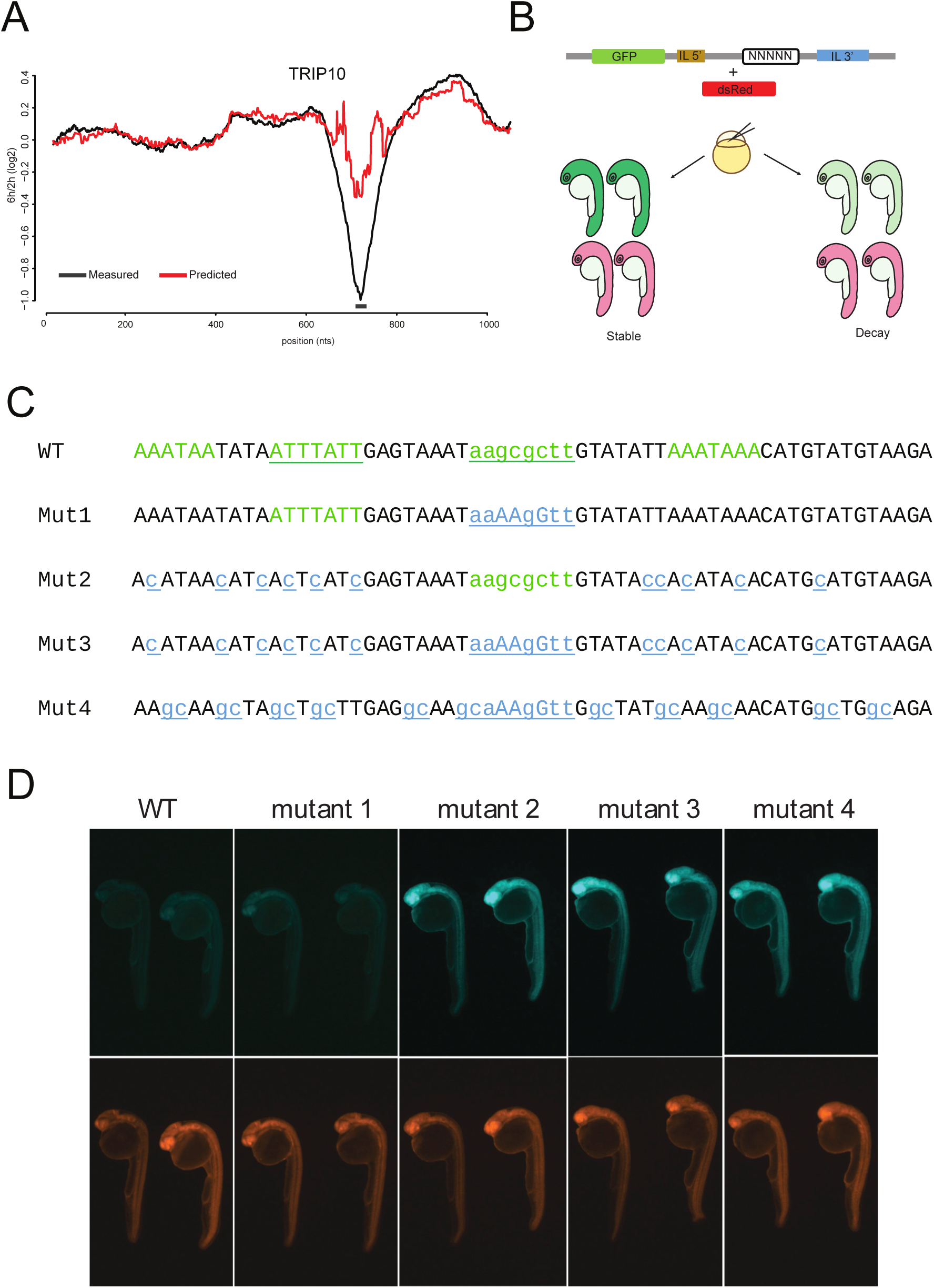
*Trip10* transcript case study. (A) RESA profile from the *trip10* locus comparing Late/early fold change (black curve) and predicted *trip10* profile according to the RF model (red curve) (B) experimental design for validating the activity of injected mRNA reporters by comparing GPF to the control dsRed. (C) mRNA reporter sequence with miR-430 motif with a GU wobble and multiple AUUUA sequences motifs (green), while blue color represents the mutation introduced for the design of each validation reporter. (D) Assessment of each injected mRNA reporter effect on stability. Comparing the lower control panel (red), and the GFP intensity which corresponds to stability.

**Figure S13.**
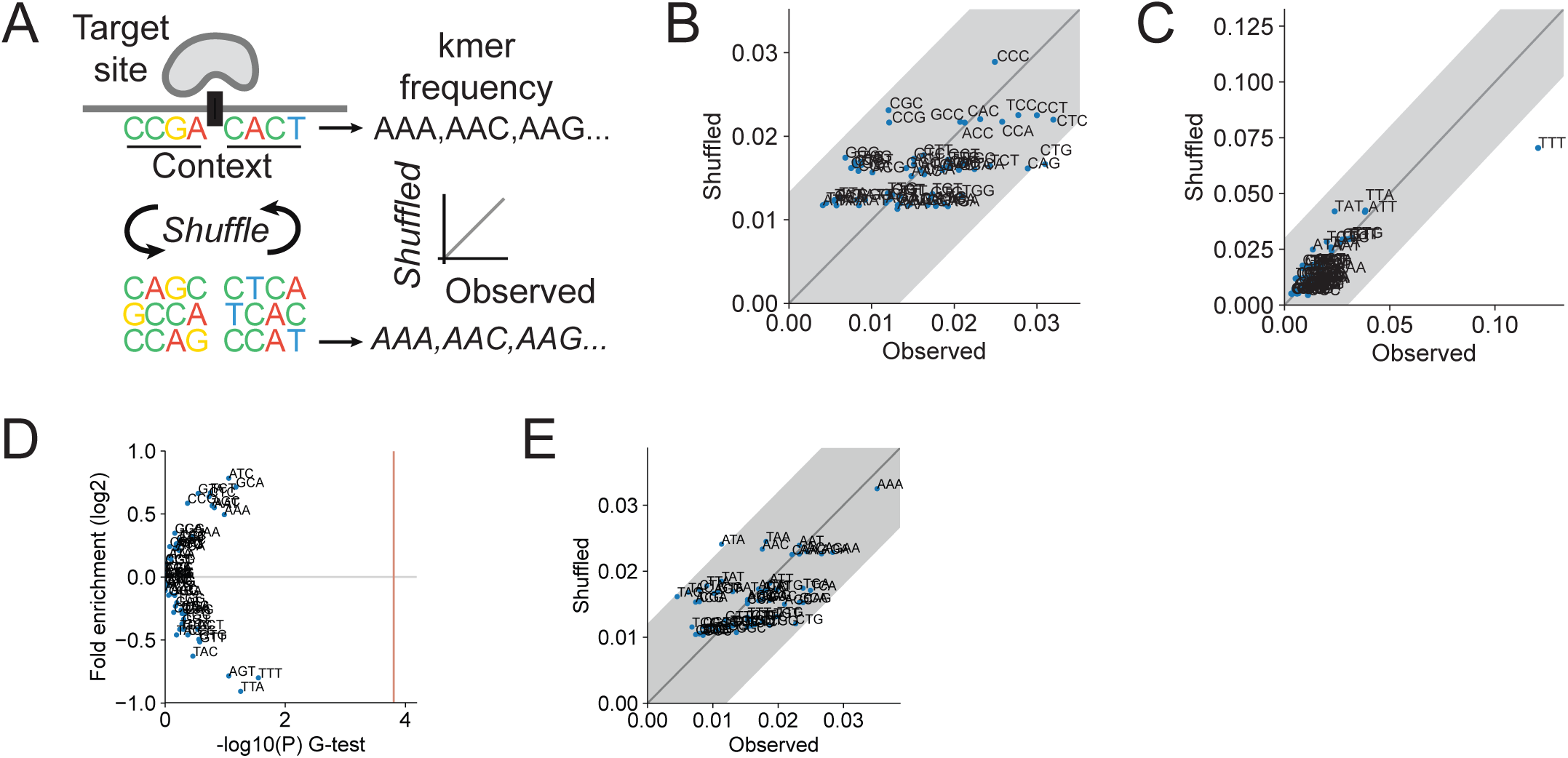
Dependency between nucleotide within target context. (A) Diagram showing the strategy used to determine if the context was preferentially enriched with specific motifs or limited to single nucleotide. Context sequences were shuffled keeping the same nucleotide composition. (B) Shown is similar motif frequency between observed and shuffled context, suggesting that the key feature of *pcbp2* binding context is nucleotide frequency. (C) For *hug*, the UUU 3mer was enriched flanking *hug* binding sites compared to shuffled sequences, demonstrating a preference for adjacent Us in favorable *hug* binding conferring stabilization. (D) Volcano plot representing 3-mer enrichment 20 nt upstream and downstream motif between the top 10% most destabilizing for miR-430 and the bottom 10%. P-values were calculated using G-test. Red line indicates 1% significance cutoff after Bonferroni multiple test correction. (E) Same as (B) for miR-430.

